# Enrichment experience improves hippocampal sparse coding via inhibitory circuit plasticity

**DOI:** 10.64898/2026.03.31.715605

**Authors:** Ekaterina Verdiyan, Stylianos Kouvaros, Joana I. Gomes, Josef Bischofberger

## Abstract

Environmental enrichment enhances hippocampus-dependent learning and memory, yet the underlying circuit mechanisms remain largely unknown. Here we combined miniscope calcium imaging during spatial exploration with synaptic circuit analysis in hippocampal slices to determine how enrichment experience alters hippocampal network dynamics. Prolonged enrichment reduced average firing rates and immediate early gene expression in CA1 pyramidal cells, while increasing peak firing and spatial selectivity. Population activity was sparser and more diverse, resulting in a higher Gini index. Circuit analysis revealed enhanced excitatory drive onto both pyramidal cells and somatostatin (SOM) interneurons, together with a strengthened SOM-mediated feedback inhibition onto pyramidal cells. Suppressing SOM interneurons occluded the enrichment-induced augmentation of sparsity and Gini index and prevented improvements in hippocampus-dependent learning. These findings demonstrate that environmental enrichment dynamically enhances hippocampal sparse coding through potentiation of SOM-mediated feedback inhibition, linking experience-dependent inhibitory plasticity to enhanced memory performance.

## Introduction

Environmental enrichment (EE) has been intensively investigated for its potential to enhance cognitive activity and improve learning and memory^1,2^. The interest in this topic is steadily increasing, because enrichment represents a promising lifestyle intervention for mitigating cognitive decline in aging and brain disorders^3–6^. Nevertheless, the circuit-level mechanisms underlying enrichment-dependent improvements in brain function remain still poorly understood.

At the molecular level, hundreds of genes exhibit differential expression following EE, which generally seems to promote more juvenile-like transcriptional expression patterns in aged animals^7,8^. Furthermore, glutamatergic synapses have been extensively studied in the context of enrichment, typically showing enhanced synaptic plasticity and increased spine density, which would imply more excitation^9–12^. However, no overall increase in activity is observed at the network level. By contrast, several studies of hippocampal function report reduced activation of hippocampal principal cells in enriched animals^13,14^. Indeed, sparse coding is widely considered to be critically important for hippocampus-dependent declarative learning and memory in both animals and humans^15–20^. Specifically, confining high activity to only a small fraction of neurons may be essential for processing high dimensional information, for encoding new related memories and for building up a complex cognitive ‘knowledge structure’^21–23^.

Sparse coding critically depends on inhibitory interneurons^15,24,25^. Moreover, interneurons are dynamically modulated during novelty recognition and learning^26,27^, and inhibitory synapses are remodelled during learning to stabilize neuronal network function at multiple levels, including excitation-inhibition (E-I) balance, response normalization and gating of excitatory plasticity^28,29^. Most importantly, neuronal inhibition may serve to protect stored memories from interference during continuous learning^30^.

Among the different types of interneurons, dendrite-targeting interneurons exert particularly powerful inhibitory control, as they regulate dendritic NMDA electrogenesis and synaptic plasticity in pyramidal cells (PCs)^31–34^. In addition, they shape synaptic integration and govern burst firing^35,36^. Most importantly, both the size of neuronal cell assemblies and the precision of learning is dependent on somatostatin-positive (SOM) dendrite-targeting interneurons^37–39^. Despite their central role in learning and memory, the impact of enrichment experience on interneuron function remains largely unknown.

To understand the role of interneurons in EE, we focused on dendrite-targeting hippocampal SOM interneurons in the CA1 region. By combining rigorous circuit analysis in acute brain slices with optogenetics and calcium imaging in freely moving mice, we identified a major contribution of SOM interneurons to improved sparse coding and enhanced hippocampal learning and memory following environmental enrichment experience.

## Results

### Enrichment improves sparsity and spatial selectivity of pyramidal cell firing

Neuronal circuit mechanisms in EE were examined in adult mice, which were housed either in standard cages (STD) or in larger rat cages containing a variety of objects and a running wheel for several weeks (EE, Supplementary Fig. 1). To investigate activity of CA1 pyramidal cells in freely moving mice, we performed *in vivo* calcium imaging using an AAV.CaMKII.GCaMP6f virus, in combination with an implanted gradient-index (GRIN) prism lens and a miniature microscope (Fig. 1a-c). Animals were habituated to both the microscope and the recording arena (80 x 80cm), containing fixed objects for more than a week. Following habituation, mice were placed into the arena and calcium-dependent fluorescence transients were recorded during 15-min exploration sessions. Somatic fluorescence traces were extracted and the relative fluorescence change was calculated as df = Δf/f_0_ (Fig. 1d-e).

**Figure 1.**
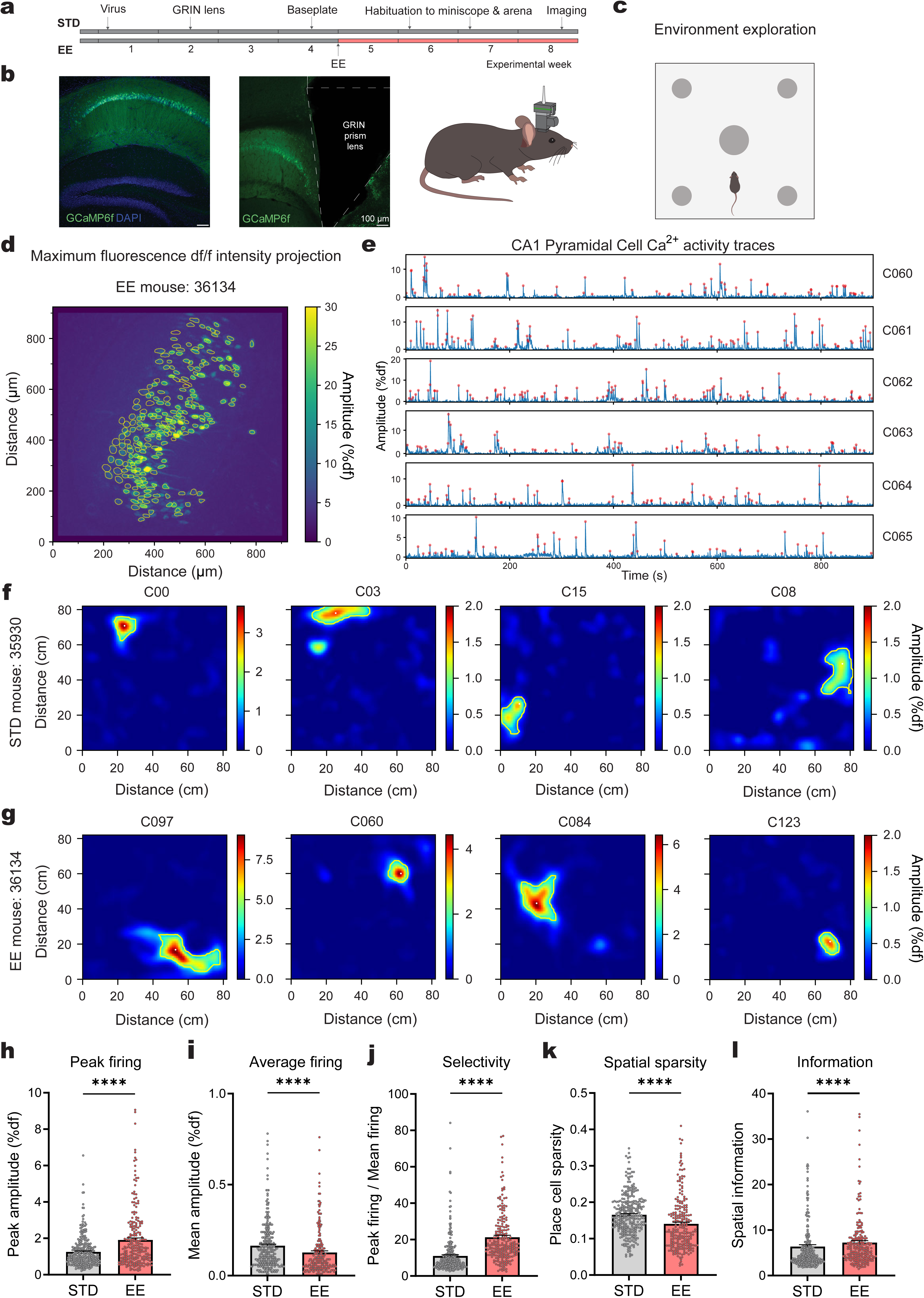
Enrichment enhances spatial selectivity of CA1 pyramidal cells during spatial exploration. **a)** Experimental timeline. **b)** Confocal images showing CaMKII-GCaMP6f expression in CA1 pyramidal cells (left) and location of GRIN prism lens within dorsal CA1 (right). – GCaMP6f – green, DAPI – blue. Scale bar, 100 µm. **c)** Open field arena (80 x 80 cm) for environmental exploration and recording of Ca^2+^ activity in CA1 PCs (455 nm LED). Duration of a recording session: 15 min. **d)** Representative image of a maximum df/f intensity projection from dorsal CA1 recorded during a 15-min spatial exploration. The image shows all cells that were active throughout the session. Cell bodies of PCs are outlined as ROIs, which were automatically identified using PCA-ICA and used for extraction of individual fluorescence traces from recorded movies. Axes scaling: 1.20 µm/pixel. **e)** Example fluorescence Ca^2+^ activity traces from 6 CA1 PCs during 15-min exploration. Identified events are marked with red dots. **f-g)** Spatial firing maps for 4 PCs from one example STD mouse (35930) (**f**) and for 4 PCs from one example EE mouse (36134) (**g**). **h-l)** Bar graphs showing (**h**): Higher peak amplitude of calcium within place field (p<0.0001, Mann-Whitney test), (**i**): Lower mean firing amplitude across the entire arena in place cells (p<0.0001, Mann-Whitney test), (**j**): Increased place cell spatial selectivity (p<0.0001, Mann-Whitney test), (**k**): Enhanced place cell sparsity (p<0.0001, Mann-Whitney test), and (**l**): Higher spatial information per place cell in EE compared to STD mice (p<0.0001, Mann-Whitney test). Number of cells for all bar graphs are n=289 cells, N=4 mice for EE and n=230 cells, N=4 mice for STD. Dots represent cells. Data are shown as mean ± SEM. **** corresponds to p<0.0001.

We then derived spatial firing fields by averaging Ca^2+^ activity for each cell within small spatial bins (1.2 x 1.2 cm), showing typical place-field firing in a similar proportion of recorded pyramidal cells from standard (289 of 440 cells, N=4) and enriched animals (230 of 464 cells, N=4, Fig. 1f-g). Quantitative analysis revealed a higher peak Ca^2+^ amplitude in enriched mice at the center of the place fields (1.91 ± 0.11 %df vs 1.25 ± 0.05 %df in n=230 and n=289 cells, p<0.0001, Fig. 1h). Due to lower average Ca^2+^ activity across the entire arena in enriched mice (Fig. 1i), the spatial selectivity - calculated as peak over mean activity - was approximately twofold larger in mice with enrichment experience (21.3 ± 1.1 vs 11.0 ± 0.6, p<0.0001, Fig. 1j). Furthermore, the overall sparsity was more pronounced (Fig. 1k) and spatial information per place cell was increased in enriched mice (7.25 ± 0.44 vs 6.38 ± 0.41, p<0.0001, Fig. 1l).

Analysis of firing across all PCs revealed a lower frequency of Ca^2+^ events in enriched compared to standard mice (0.033 ± 0.001Hz, n=464 vs 0.051 ± 0.001 Hz, n=440, p<0.0001, Fig. 2a). Furthermore, we quantified the proportion of co-active cells within 10-s time bins (Fig. 2c), which was also reduced (19.9 ± 0.6% vs 31.0 ± 0.8%, n=360 bins each, p<0.0001). Interestingly, the mean event amplitude per cell, averaged throughout the 15-min session, was not different between groups (p=0.6102, Fig. 2b). However, the cumulative distribution of the event amplitudes across cells exhibited a smaller slope, indicating that the activity is more unequally distributed between different neurons. Given that pyramidal cell firing follows a logarithmically skewed distribution^21^, we quantified the firing activity as the product of event frequency and amplitude in individual cells and performed a logarithmic transformation (Fig. 2d). The standard deviation of a fitted gaussian function to the log-transformed distribution was 1.4-times broader in enriched mice (σ_EE_ = 0.426 vs σ_STD_ = 0.301, F_(1,24)_ = 22.34, p<0.0001). Similarly, the distribution of mean amplitudes in different neurons was 1.6-fold broader in enrichment compared to standard housing (σ_EE_ = 0.237 vs σ_STD_ = 0.149, F_(1,36)_ = 60.78, p<0.0001, Fig. 2e).

**Figure 2.**
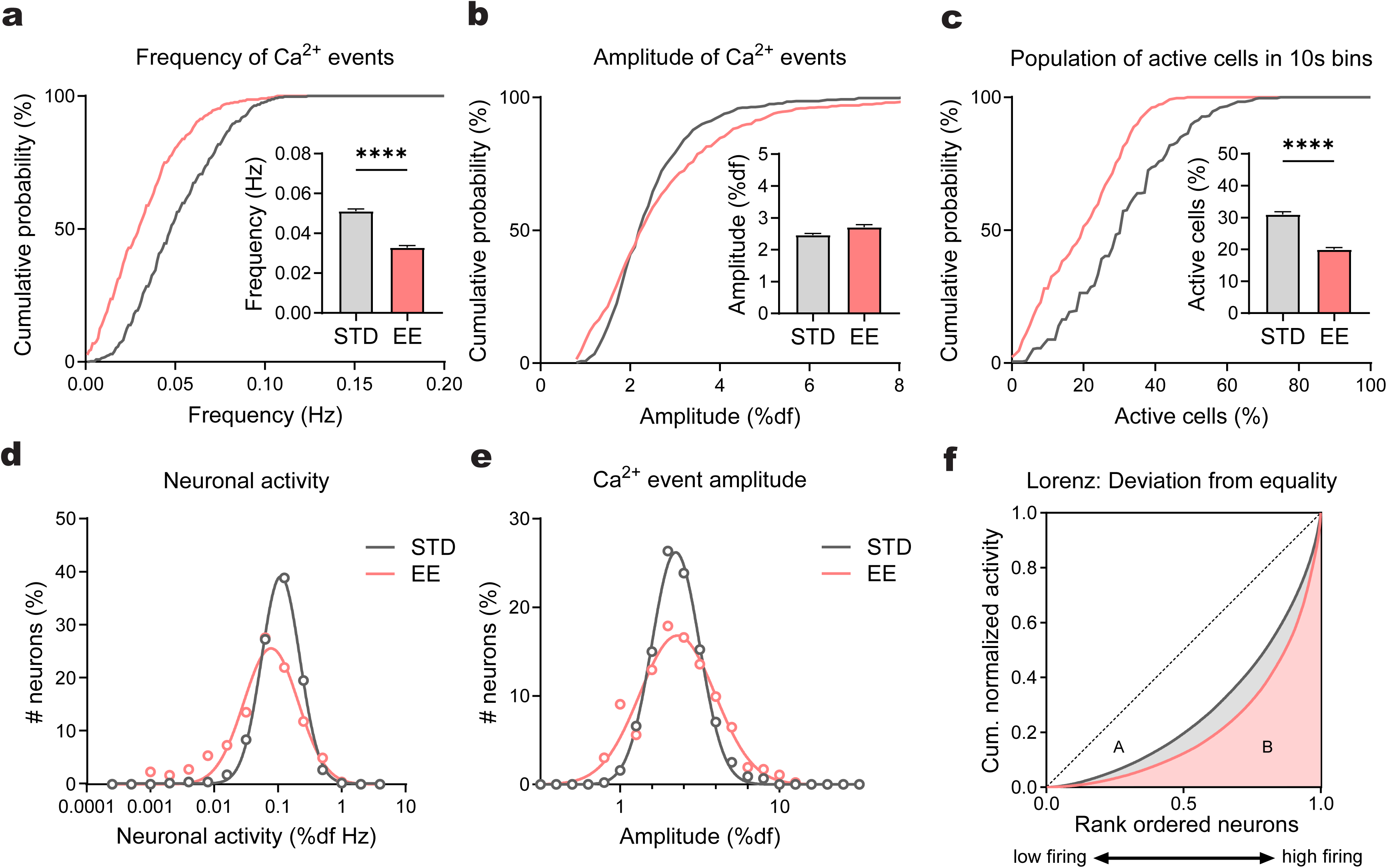
Enrichment increases sparsity and diversity of CA1 pyramidal cell firing. **a)** Cumulative distribution shows reduced mean Ca^2+^ event frequency in CA1 PCs of EE compared to STD (p<0.0001, Kolmogorov-Smirnov test). Inset: Reduced average firing frequency in EE (p<0.0001, Mann-Whitney test). EE: n=464, N=4 vs STD: n=440, N=4. **b)** Cumulative distribution of mean Ca^2+^ event amplitudes in CA1 PCs in EE compared to STD housing (p<0.0001, Kolmogorov-Smirnov test). Inset: average amplitudes in EE and STD mice (p=0.6102, Mann-Whitney test). Same cells as in (a). **c)** Cumulative distribution shows a smaller number of co-active PCs within the same 10-s time bin in EE mice. (p<0.0001, Kolmogorov-Smirnov test). Inset: Reduced average number of co-active CA1 PCs in EE mice (p<0.0001, Mann-Whitney test). Same cells as in (a). **d)** Broader log-transformed distribution of neuronal activity, quantified as the product of event frequency and amplitude, in EE (F_(1,24)_ = 22.34, p<0.0001). Same cells as in (a). **e)** Broader log-transformed distribution of mean Ca^2+^ event amplitudes in EE compared to STD (F_(1,36)_ = 60.78, p<0.0001). Same cells as in (a). **f)** Lorenz curve, showing a more unequal cumulative distribution of firing activity across rank-ordered population of all active cells in EE mice. Equal firing would be represented by the identity line (dashed line). Gini index, calculated as deviation from linear growth equal to the area below Lorenz curve (B) normalized to the area below the identity line (A+B), is larger in EE (red) compared to STD mice (grey, p<0.0001). Same cells as in (a). Data are shown as mean ± SEM. **** corresponds to p<0.0001.

Finally, we quantified the diversity and inequality of firing across different neurons using a Lorenz curve^24^. For this analysis, the cumulative activity of cells was plotted against the rank-ordered number of neurons. In case all neurons would fire with the same mean frequency, this graph would follow an identity line (Fig. 2f, dotted line). However, the real data are distributed in a highly non-linear fashion, exceeding 50% population activity with as many as 79.7% neurons in STD (gray line) and 87.3% of neurons in EE mice (red line). This shows that only a relatively small number of cells of 20% and 13% contribute half of the overall activity in STD and EE mice, respectively. This unequal distribution can be further quantified as deviation from linear growth (area A), normalized to the total area below the identity line (A+B). This value ranging from 0-1 is called Gini index and is larger in enriched mice (0.58 versus 0.45, p<0.0001).

These data demonstrate that hippocampal information processing during spatial exploration differs markedly in animals with enrichment experience. Only 13% of the active cells account for approximately half of the overall activity, whereas a substantially larger fraction of highly active cells contributes to population activity in STD housing. Although the proportion of place cells appears to be similar between groups, place cell activity in enrichment is characterized by higher peak firing within place fields and increased spatial selectivity, with lower firing rates outside place fields and higher spatial information per cell.

### Reduced cFos expression in enriched mice during spatial exploration

Place field formation in CA1 pyramidal cells has been shown to depend on the expression of the immediate early gene cFos^40^. Therefore, we investigated cFos expression during exploration of a novel environment (NE) using immunohistochemistry. Mice were either taken directly from home cage or sacrificed and perfused after 1h of exploration in a novel arena containing novel objects at different locations (Supplementary Fig. 1c). As shown in Fig. 3a-d, NE exploration induced robust cFos expression in a substantial fraction of neurons within the CA1 pyramidal cell layer in both groups. However, the proportion of cFos-positive neurons was lower in mice after 6 weeks of EE compared to STD mice (8.9% ± 0.7% vs 14.7% ± 0.7%, N=11 mice each, p<0.0001, Fig. 3e). A similar reduction of exploration-induced cFos expression was already apparent after 2 weeks of enrichment (9.0% ± 0.9% vs 14.7% ± 0.7%, N=7 and N=11, p=0.0204, Fig. 3f). In addition to the reduced number of cFos-positive cells in enriched mice, the mean fluorescence intensity per labeled cell was also lower (p<0.0001, Fig. 3g-h).

**Figure 3.**
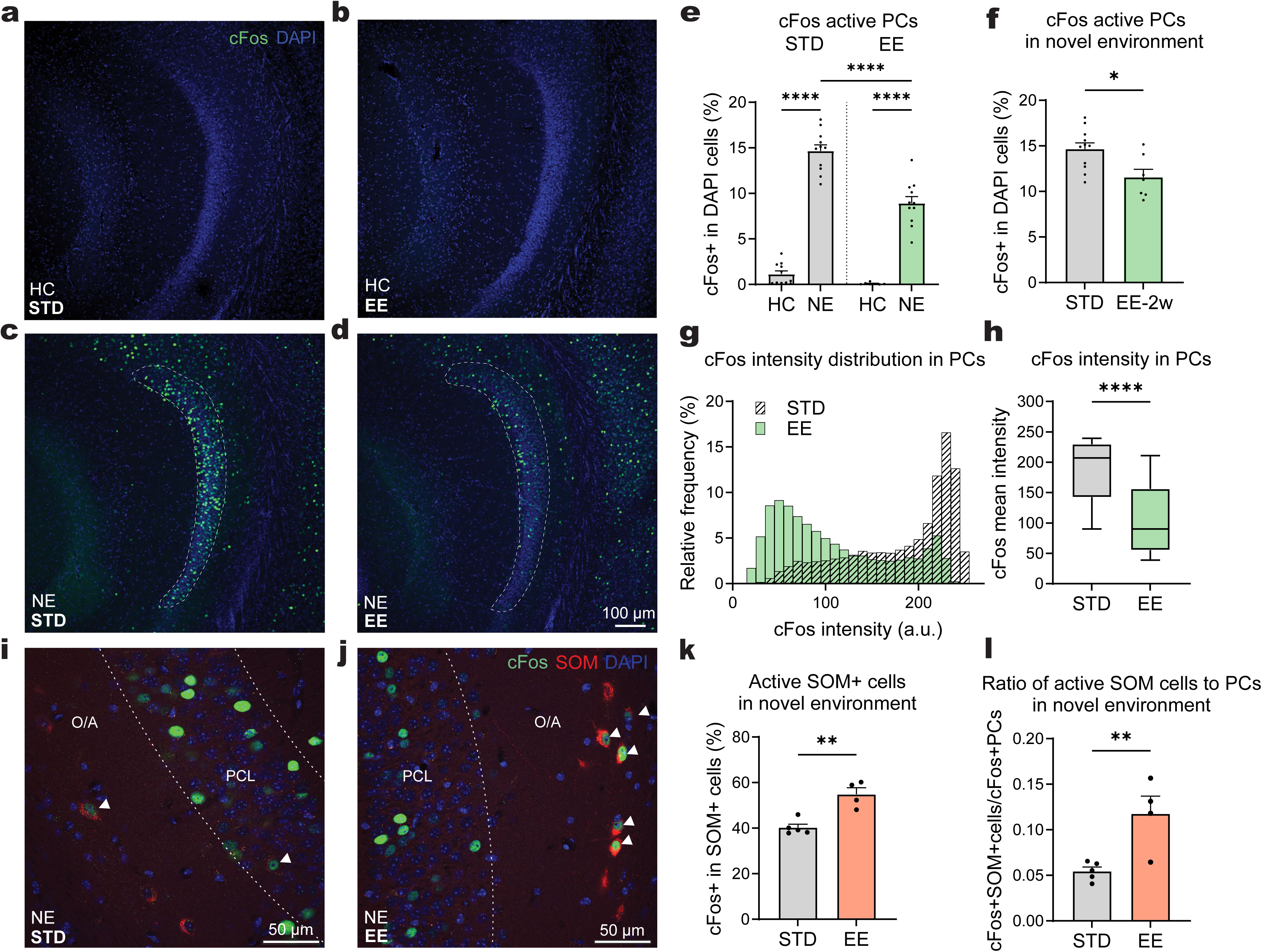
Opposing impact of enrichment on cFos activity in CA1 PCs and SOM interneurons. **a-d)** Representative confocal images showing cFos expression in dorsal CA1 of a STD mouse (**a**, **c**) and an EE mouse (**b**, **d**). While cFos is low in animals from home cage (HC, **a-b**), its activity strongly increases during spatial exploration (1h) of novel environment (NE) in STD mice (**c**). A substantially smaller increase was observed in EE mice (**d**). cFos - green, DAPI - blue. Scale bar, 100 μm. **e)** Reduced cFos expression in CA1 PCs in EE as compared to STD mice after NE exploration (EE-NE vs STD-NE: p<0.0001, two-way ANOVA with Tukey’s multiple comparisons test; EE: N=7 and N=11, STD: N=11 and N=11, for HC and NE, respectively). **f)** Comparison of cFos active cells after exploration in STD housing (same data as in **e**) vs 2 weeks of EE (p=0.0126, unpaired two-sided t-test, STD: N=11, EE-2w: N=7). **g)** Histogram of distribution of cFos intensity in CA1 PCs after spatial exploration in STD housing (n=26617 cells, N=10) vs. EE mice (n=16634 cells, N=11). **h)** cFos intensity in CA1 PCs is lower in EE mice (p<0.0001, Mann-Whitney test; same cells as in **g**). Box plots represent median (horizontal line) and percentile ranges (25th-75th, box; 10th-90th, whiskers). **i- j)** Representative confocal images showing cFos expression in SOM interneurons (cFos^+^ SOM^+^, white arrowheads) in dorsal CA1 of mice from the STD (**i**) and EE (**j**) after NE exploration. Abbreviations: PCL - pyramidal cell layer, O/A – oriens/alveus. cFos - green, SOM - red, DAPI - blue. Scale bar, 50 μm. **k)** Increased cFos activity in SOM interneurons (cFos+SOM+ / SOM+ cells) in EE compared to STD after spatial exploration (p=0.0019, unpaired t-test, EE: N=4 vs STD: N=5). **l)** Higher ratio of cFos+SOM+ interneurons to cFos+PCs in enriched mice after spatial exploration compared to STD mice (p=0.0171, unpaired t-test, same animals as in **k**). Dots represent individual mice. Data are shown as mean ± SEM. * p<0.05, ** p<0.01 and **** p<0.0001.

Because SOM interneurons were suggested to regulate the size of hippocampal cell assemblies^38^, we asked whether their activation is altered by enrichment conditions. Indeed, cFos expression in somatostatin-positive interneurons was higher in EE mice (54.9% ± 2.8% vs 40.3% ± 1.5%, N=4 and N=5, p=0.0019, Fig. 3k). As this pattern contrasted with the reduced activation of PCs, we calculated the ratio of active SOM interneurons to active PCs. This ratio was 2-times larger in EE (11.5% ± 2.1% vs 5.5% ± 0.4%, p=0.0171, Fig. 3l), indicating that SOM interneurons are 2-fold more likely to be recruited during spatial exploration in enriched mice. Together, these findings suggest that enrichment experience promotes sparser activation of CA1 pyramidal cells, accompanied by enhanced recruitment of SOM interneurons. This cell-type specific shift in activation implies that enrichment alters hippocampal circuit dynamics and modifies hippocampal information processing.

### Increased glutamatergic excitation of SOM interneurons

Reduced activation of CA1 pyramidal cells in EE could be due to decreased glutamatergic excitation or due to enhanced GABAergic inhibition, potentially via SOM interneurons. To examine synaptic connectivity, we performed whole-cell patch-clamp recordings in acute hippocampal slices. As a first step, we stimulated Schaffer collaterals and recorded evoked excitatory postsynaptic currents (EPSCs) in PCs. Notably, the maximal EPSC amplitude was approximately 2-fold larger in PCs of EE mice (744 ± 117 pA vs 398 ± 80 pA, n=14 and n=11 cells, p=0.0301, Supplementary Fig. 2). Furthermore, the frequency of miniature EPSCs (mEPSCs) was approximately 2-fold higher in EE mice compared to STD (3.3 ± 0.8 Hz vs 1.5 ± 0.2 Hz, n=13 each, p=0.0042, Supplementary Fig. 2). Therefore, the number of glutamatergic synapses in pyramidal cells is not smaller but actually 2-fold larger in enriched mice.

Next, we asked whether the density of glutamatergic synapses onto interneurons might also be altered in EE. To address this question, we stained for glutamatergic synapses in SOM interneurons using a recently developed Cre-dependent Fibronectin intrabodies generated by mRNA display (FingRs) directed against PSD95, selectively expressed in SOM interneurons of SOM-Cre-tdTomato mice^41,42^ (Fig. 4a). The FingR construct was fused to a CCR5 zinc finger domain (ZnF), mGreenLantern and a CCR5-KRAB repressor domain to keep unspecific cytosolic fluorescence to a minimum. As shown in Figure 4b-c, the density of FingR.PSD95-positive puncta is higher in SOM interneurons from EE compared to STD mice (2.10 ± 0.05 vs 1.47 ± 0.08 puncta per µm, n=63 and n=45, p<0.0001, Fig. 4c), while the size of puncta was similar (p=0.7311, Fig. 4d). These findings suggest enhanced glutamatergic synaptic input onto SOM interneurons under enrichment conditions.

**Figure 4.**
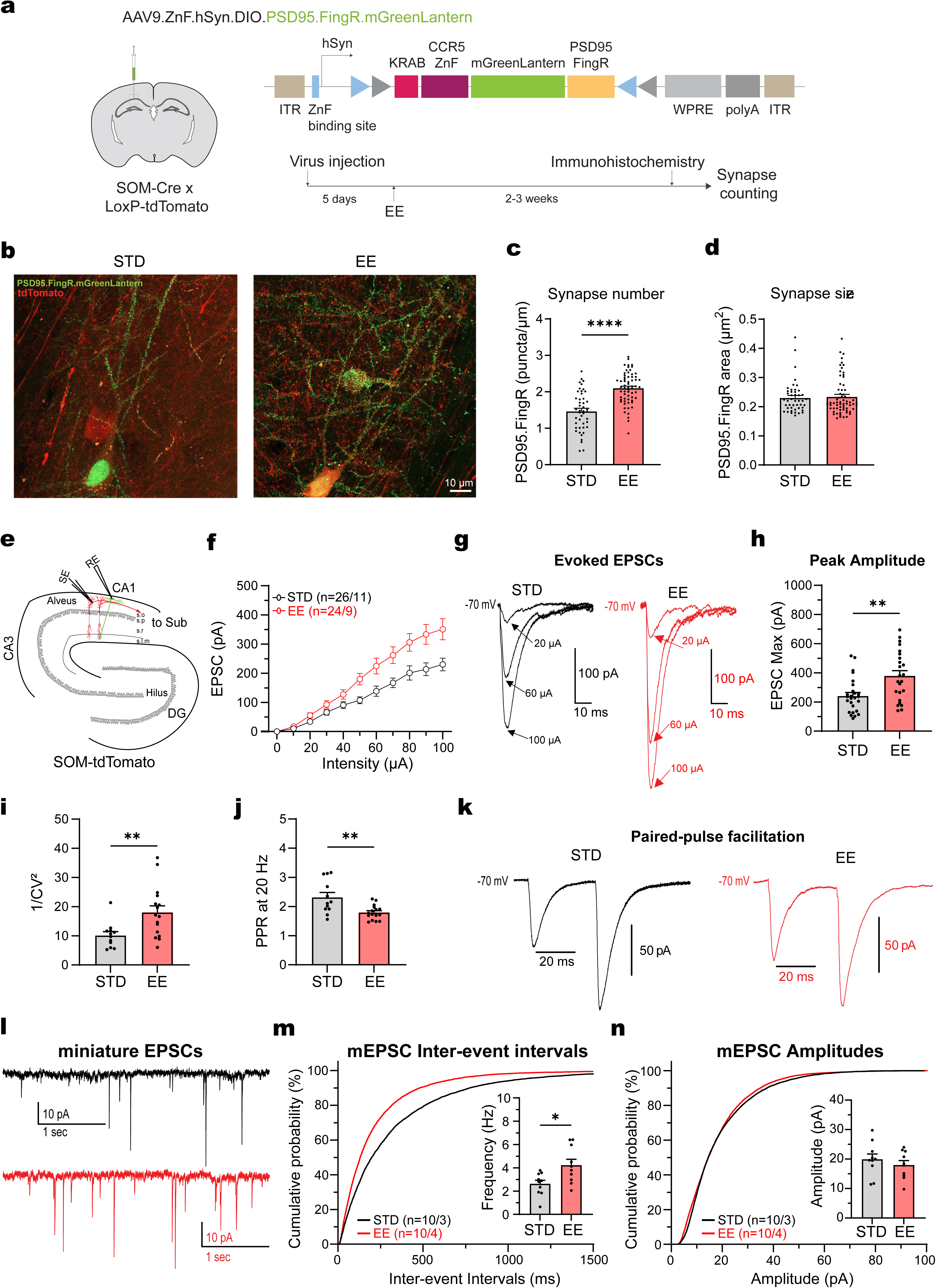
Enrichment increases excitatory drive onto CA1 SOM interneurons. **a)** Top: Schematic illustration showing AAV-driven Cre-dependent FingR intrabody probes for labelling glutamatergic synapses (PSD95.FingR-mGreenLantern). Bottom: Experimental timeline: Five days after virus injection, mice were placed into EE for 2-3 weeks. After that, immunohistochemistry and synapse counting was performed. **b)** Representative confocal images of CA1 SOM interneurons (tdTomato+, red) and glutamatergic synapses (PSD95.FingR-mGreenLantern, green) in STD (left) and EE mice (right). Scale bar, 10 μm. **c)** Higher glutamatergic synapse density on dendrites of CA1 SOM interneurons in EE compared to STD housing (p<0.0001, Mann-Whitney test, EE: n=63 vs STD, n=45 dendritic branches). **d)** No difference in the size of glutamatergic synapses between two groups (p=0.7311, Mann-Whitney test). **e)** Schematic illustration showing the recorded CA1 SOM cell along with the afferent stimulation in oriens alveus of CA1, using SOM-tdTomato mice. **f)** Stimulus-response curve showing larger evoked EPSCs in response to electrical stimulation of CA1 PC afferents in CA1 SOM cells of EE mice (red, n=24, N=9) compared to STD housing (black, n=26, N=11). Stimulation intensities were ranging from 0 to 100 μA (p=0.0001, 2-way ANOVA, F _(10, 523)_ = 57.92). **g)** Representative traces of EPSCs in CA1 SOMs evoked by electrical stimulation at different intensities of 20, 60 and 100 μA, recorded at -70 mV in presence of gabazine. **h)** Increased maximal amplitudes of evoked EPSCs in CA1 SOMs upon stimulation of CA1 PCs in EE versus STD (p=0.0024, Mann-Whitney test). **i)** Increased reliability of synaptic currents quantified as 1/CV² in EE versus STD mice (EE: 18.0 ± 2.2 vs STD: 10.2 ± 1.3, p=0.0083, Mann-Whitney test, EE: n=16, N=9 vs STD: n=12, N=7). **j)** Decreased PPR recorded at 20Hz in EE versus STD mice (EE: 1.80 ± 0.06 vs STD: 2.32 ± 0.16, p=0.0044, Mann-Whitney test, EE: n=16, N=9 vs STD: n=12, N=7). **k)** Representative traces of two consecutive EPSCs at 20 Hz in SOMs evoked by electrical stimulation of PC afferents for STD (black) and EE (red) mice. **l)** Representative traces of mEPSCs from STD (black) and EE (red) mice recorded at -70 mV in presence of gabazine and TTX. **m)** Cumulative frequency distributions showing reduced interevent intervals of mEPSCs recorded from EE SOM cells (red, p<0.0001, Kruskal-Wallis test). Inset: Increased average mEPSC frequencies per cell (p=0.0129, Unpaired t-Test, EE: n=10, N=4 vs STD: n=10, N=3). **n)** Cumulative distribution of amplitudes of mEPSCs recorded from STD (black) and EE SOM cells (red). Inset: Similar average mEPSC amplitudes per SOM cell in EE and STD mice (p=0.631, Mann-Whitney test, EE: n=10 vs STD: n=10). Data are presented as mean ± SEM. * p<0.05, ** p<0.01 and **** p<0.0001.

To assess the functional properties of these synapses, we recorded evoked EPSCs in CA1 SOM interneurons in acute hippocampal slices of SOM-tdTomato mice while stimulating PC axons (Fig. 4e-k). The amplitude of synaptic currents was larger in SOM interneurons of EE mice (380 ± 35pA vs 242 ± 23 pA, n=24 and n=26 cells, p=0.0024, Fig. 4h). Additionally, synaptic reliability quantified as 1/CV^2^ was increased and paired-pulse ratio was reduced, consistent with an elevated presynaptic release probability (Fig. 4i-k). Finally, we also analyzed mEPSCs in SOM interneurons, showing a higher event frequency in EE mice (4.24 ± 0.50 Hz vs 2.63 ± 0.30 Hz, n=10 each, p=0.0129), whereas amplitudes were similar between groups (p=0.631, Fig. 4l-n). Similar results were obtained by analyzing spontaneous EPSCs (Supplementary Fig. 3). These findings show that the density of glutamatergic synapses onto SOM interneurons is substantially higher in enrichment, resulting in approximately 2-times bigger excitatory synaptic currents.

Taken together, the data demonstrate that, besides more synapses onto CA1 pyramidal cells, enrichment experience also enhances glutamatergic connectivity onto SOM interneurons. In addition to the higher synapse number, the increased release probability further supports the conclusion that excitatory drive onto SOM interneurons is more efficient in EE compared to mice in STD housing.

### Increased SOM-interneuron mediated inhibition of CA1 pyramidal cells

To examine the inhibitory output of SOM interneurons, we injected a Cre-dependent ChR2-virus into SOM-Cre animals and recorded light-evoked inhibitory postsynaptic currents (IPSCs) in CA1 PCs (Fig. 5a). Blue light pulses (473 nm) were applied to a relatively small area (d = 50 µm) in stratum lacunosum moleculare to investigate dendritic inhibition of the recorded PCs. The maximal IPSC amplitude was approximately 2-fold larger in EE mice compared to STD (556 ± 57 pA vs 336 ± 21 pA, n=13 and n=16, p=0.0010, Fig. 5b-d). Similarly, the inhibitory charge transfer was approximately 2-fold larger (37.4 ± 2.9 pC vs 20.7 ± 1.5 pC, p<0.0001, Fig. 5e). Also, the decay time-course of IPSCs was slower in EE (57.0 ± 3.3 ms vs 45.0 ± 1.8 ms, p=0.0116, Fig. 5f), suggesting a more powerful dendritic inhibition generated by SOM interneurons. Presynaptic properties, assessed by paired-pulse ratio and coefficient of variation analysis, were not different between groups (Supplementary Fig. 4).

**Figure 5.**
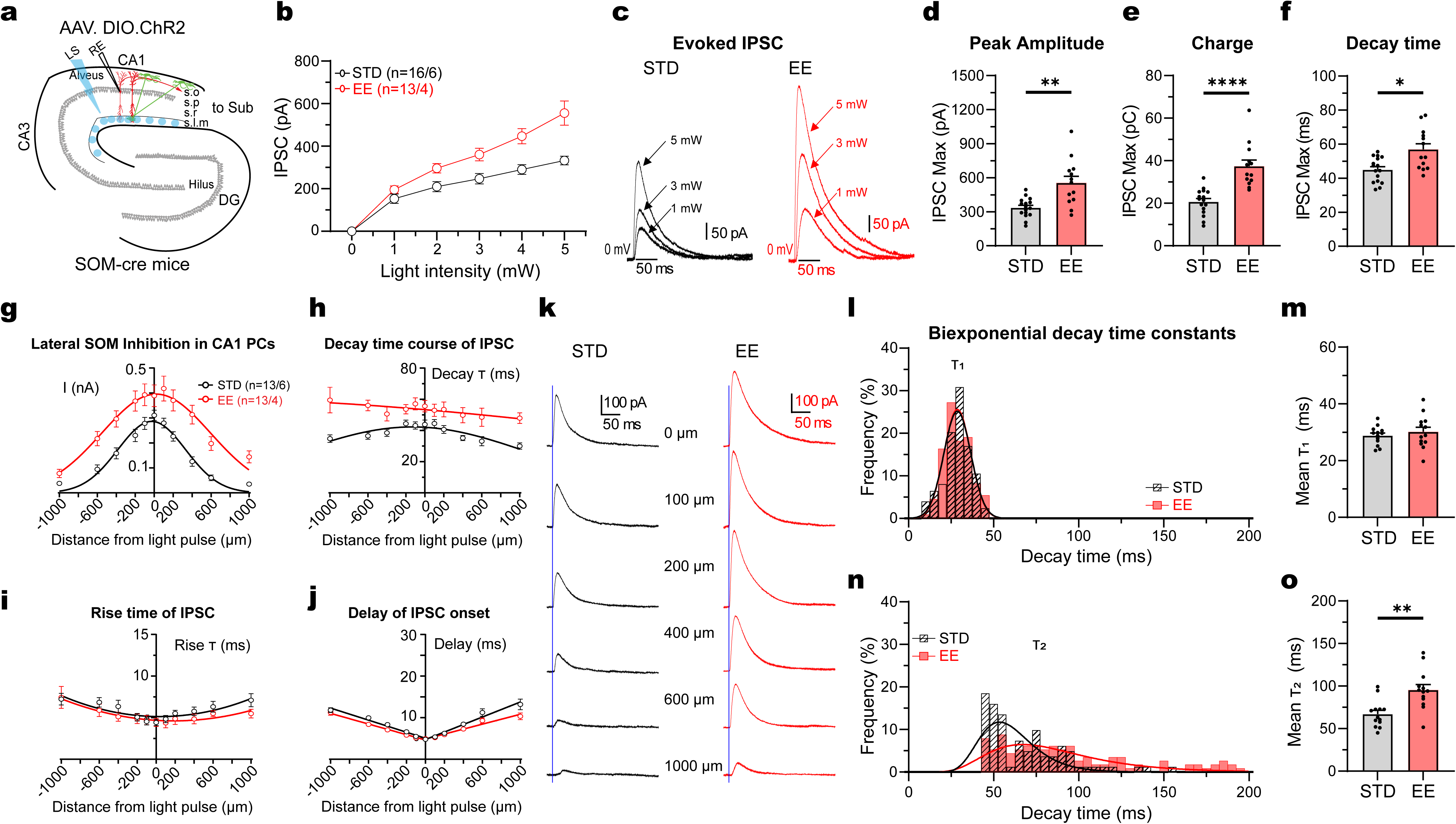
Increased inhibitory output of SOM interneurons onto CA1 PCs. **a)** Schematic illustration showing optogenetic light stimulation in stratum lacunosum moleculare (SLM) at various distances from the recorded CA1 PC, with conditional expression of AAV.DIO.ChR2 in SOM-Cre mice (+1000 μm corresponds to site towards CA3 and -1000 μm towards the subiculum). **b)** Stimulus-response curve showing larger evoked IPSCs in CA1 PCs in response to optogenetic stimulation of CA1 SOMs from EE mice (n=13, N=4) compared to STD housing (n=16, N=6) (p<0.0001, 2-way ANOVA, F _(5, 162)_ = 46.48). Stimulation intensity range: 0-5 mW. **c)** Representative traces of SOM-mediated IPSCs in CA1 PCs at different light intensities (1, 3, 5 mW) for STD (black) and EE mice (red). **d)** Larger evoked maximal SOM-mediated IPSC amplitudes in CA1 PCs from EE compared to STD mice (p=0.0010, Mann-Whitney test). **e)** Larger integral of maximal SOM-mediated IPSCs in CA1 from EE compared to STD mice (p<0.0001, Mann-Whitney test). **f)** Decay time course of the maximal SOM-mediated IPSCs in CA1 PCs in EE compared to STD (p=0.0116, Mann-Whitney test). **g-j)** Summary graphs of the average IPSC amplitude fitted with a Gaussian function (**g**), decay time course (**h**) and rise time (**i**) fitted with a quadratic function, and synaptic delay fitted by linear regression (**j**), after light stimulation in SLM at various lateral stimulation sites. Stimulation intensity of 4mW was used in both conditions. **k)** Representative average traces of SOM-mediated IPSCs (recorded at 0 mV) in CA1 PCs at different distances from STD (black) and EE mice (red). The blue vertical line points out the time of light stimulus, indicating the synaptic delay of IPSCs elicited after optogenetic SOM axons stimulation at the various distances. **l&n)** Distribution of the fast (**l**) and slow (**n**) decay time constants after fitting a biexponential function to SOM-mediated IPSCs from STD (black) and EE (red) mice. **m&o)** Comparable average (m) fast component of decay time course τ1 (STD: 28.8 ± 0.91 ms vs EE: 30.14 ± 1.61 ms, p=0.6498, Mann-Whitney test) but prolonged (**o**) slow decay component τ2 in EE (n=13, N=4) compared to STD mice (n=13, N=6, p=0.0015, Mann-Whitney test). Data are presented as Mean ± SEM. * p<0.05, ** p<0.01, and **** p<0.0001.

To explore the spatial extent of the SOM-mediated inhibition, we specifically investigated lateral inhibition by varying the stimulation site within a range of ±1000 µm relative to the recorded CA1 PC (Fig. 5a, g-k). The data revealed that the IPSCs were not only bigger with local stimulation, but the effect of enrichment was even more pronounced at more lateral stimulation sites. The spatial distribution of IPSC amplitudes could be fitted with a Gaussian function (Fig. 5g), which was significantly different in the two groups (p<0.0001, F_(3,270)_ = 36.15). The fitted standard deviation in EE was about 1.6-times broader than in STD mice (σ_EE_ = 573 µm vs σ_STD_ = 356 µm). This corresponds to a full width at half maximal amplitude of 1349 µm in EE compared to 838 µm in STD, suggesting that the axons of SOM interneurons in EE extend further and form synapses at distant targets more effectively.

In addition, the IPSC decay time course across the different stimulation sites was slower in EE mice (p<0.0001, F_(1,244)_ = 92.35, Fig. 5h). In contrast, IPSC rise time and latency were unchanged (p=0.4545 and p=0.9934, Fig. 5i-j). The IPSC decay time course was best described by a biexponential function fitted with the sum of two exponentials (Supplementary Fig. 5), and the values shown in Fig. 5f and Fig. 5h represent the amplitude-weighted average time constant. Fig. 5l shows a histogram of the isolated fast decay component, which could be fitted with similar Gaussian functions in EE and STD mice with a mean of 28.7ms (SD_EE_

= 7.79ms) and 28.7ms (SD_STD_ = 7.92ms), respectively (F_(1,76)_ = 0.3159, p=0.5757, n=13 and n=13, Fig. 5l-m). By contrast, the slow decay component was slower (F_(1,76)_ = 12.99, p=0.0006, Fig. 5n), showing a 1.4-times larger arithmetic mean value in EE mice (95.5 ± 6.4 ms vs 67.09 ± 4.4 ms, p=0.0015, Fig. 5o). These findings indicate that SOM interneurons generate more efficient and more widespread inhibition via larger amplitudes and prolonged decay time course of IPSCs.

Hippocampal SOM interneurons are classical feedback interneurons that mediate lateral inhibition between neighboring CA1 PCs under physiological conditions. To investigate feedback inhibition in CA1, we injected AAV.CaMKII.ChR2 into the CA1 area and recorded light-evoked IPSCs in CA1 PCs (Fig. 6a). As in the previous experiments, the illumination area was relatively small, with a diameter of 50 µm. Stimulation at a distance of 200 µm from the recorded CA1 PC evoked 2-times larger feedback inhibition in EE compared to STD mice (2150 ± 379 pA, n=11 vs 1090 ± 114 pA, n=16, p=0.0046, Fig. 6b-d). In addition, both the total inhibitory charge and the decay time constant were larger in EE mice (Fig. 6e-f).

**Figure 6.**
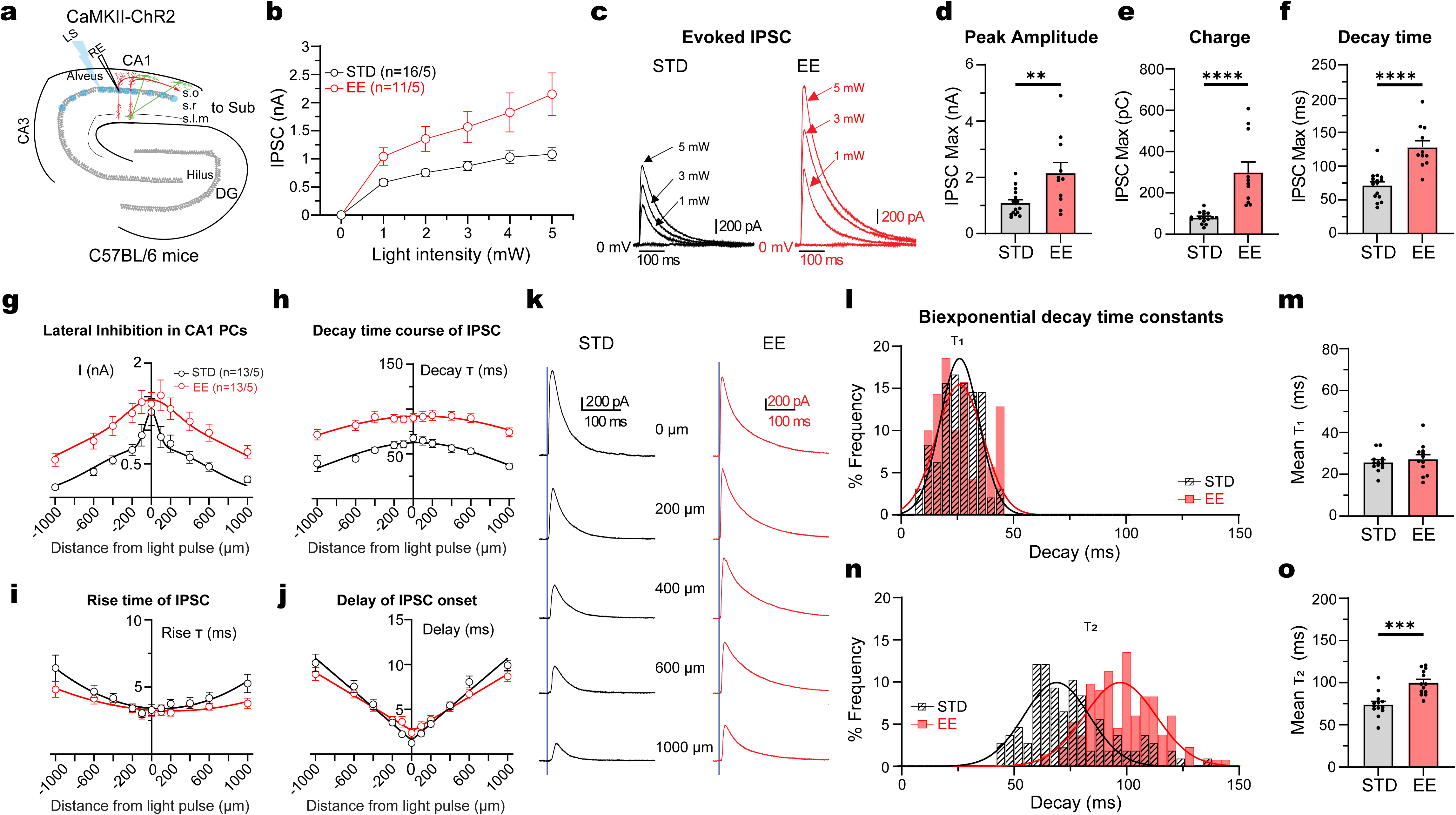
Increased lateral feedback inhibition in CA1 PCs. **a)** Schematic illustration showing optogenetic light stimulation in stratum pyramidale at various distances from the recorded CA1 PC from a mouse injected with AAV.CaMKII.ChR2 into CA1 (+1000 μm corresponds to site towards the CA3 side and -1000 μm towards the subiculum). **b)** Stimulus-response curve showing larger evoked feedback IPSCs in response to optogenetic stimulation of CA1 PCs recorded from an adjacent CA1 PC (distance 200 μm) in EE mice (p<0.0001, 2-way ANOVA, F _(1, 150)_ = 38.96, EE: n=11, N=5 vs STD: n=16, N=5). Stimulation intensity range: 0-5 mW. **c)** Representative traces of IPSCs in CA1 PCs evoked by light stimulation of adjacent CA1 at different light intensities of 1, 3 and 5 mW for STD (black) and EE (red) mice. **d)** Larger maximal evoked feedback IPSC amplitudes in CA1 PCs upon light stimulation of adjacent CA1 PCs in EE compared to STD housing (p=0.0046, unpaired t-test). **e)** Larger integral of maximal feedback IPSCs in CA1 PCs in EE compared to STD (EE: 298.4 ± 51.45pC vs STD: 81.08 ± 6.67pC, p<0.0001, Mann-Whitney test). **f)** Prolonged decay time course of maximal feedback IPSCs in CA1 PCs in EE compared to STD (EE: 127.9 ± 9.78ms vs STD: 71.3 ± 5.25ms, p<0.0001, unpaired t-test). **g-j)** Summary graphs of the average IPSC amplitude fitted with a gaussian function (g), decay time course (**h**) and rise time (**i**) fitted with a quadratic function, and synaptic delay fitted by linear regression (**j**), after light stimulation of stratum pyramidale at various distances laterally from the recorded CA1 PCs. Stimulation intensity: 3mW. **k)** Representative average traces of IPSCs (recorded at 0 mV) in CA1 PCs evoked by light stimulation of CA1 PCs at different distances for STD (black) and EE (red) mice. **l&n)** Distribution of the fast (**l**) and slow (n) decay time constants after fitting a biexponential function to feedback IPSCs from STD (black) and EE (red) mice. **m&o)** Comparable average (**m**) fast component of decay time course τ1 (p=0.5382, Mann-Whitney test) but prolonged (**o**) slow decay component τ2 in EE (n=13, N=5) compared to STD mice (n=13, N=5, p=0.0001, Mann-Whitney test. Data are presented as Mean ± SEM. ** p<0.01, *** p<0.001 and **** p<0.0001.

Next, we varied the position of the stimulating light pulse within a range of ±1000 µm relative to the recorded CA1 PC (Fig. 6g-k). The amplitude of lateral feedback inhibition was larger in EE mice, with the difference becoming more pronounced at more lateral stimulation sites (F_(1,244)_ = 64.38, p<0.0001) (Fig. 6g). Interestingly, the width of lateral feedback inhibition at half-maximal amplitude was 1404 µm in EE compared to 650 µm in STD animals, closely matching the spatial profile observed for SOM-mediated inhibitory output. We next analyzed the kinetics of IPSCs. The decay time course was fitted with a biexponential function and the amplitude-weighted time constant (τ) was calculated. As shown in Fig. 6h, the decay time course of lateral feedback inhibition was slower in EE (F_(1,239)_ = 232.1, p<0.0001), consistent with the prolonged IPSCs observed following optogenetic activation of SOM interneurons (Fig. 5h). In contrast, IPSC rise time and latency were unchanged between groups (p=0.4611 and p=0.8079, Fig. 6i, j). Separate analysis of the fast and slow components of the biexponential decay time course (Supplementary Fig. 6), revealed that the fast component was similar in both groups (27.2 ± 2.2 ms, n=13 vs 25.7 ± 1.2 ms, n=13, p=0.5382, Fig. 6l, m). By contrast, the slow component was prolonged in EE mice (99.92 ± 4.0 ms vs 74.0 ± 3.9 ms, n=13 each, p=0.0001, Fig. 6n, o).

These data show that, in addition to the enhanced glutamatergic excitation, SOM interneurons generate stronger and spatially more extended inhibitory output in enriched animals. As a consequence, these circuit properties increase lateral feedback inhibition and may therefore amplify inhibitory competition among active PCs during hippocampal information processing.

### Enhanced SOM interneuron-mediated inhibition during spatial exploration

What is the contribution of the enhanced SOM-interneuron mediated inhibition to pyramidal cell firing *in vivo*? To investigate this question, we recorded calcium activity in PCs during exploration of the same arena used above, with and without optogenetic silencing of SOM interneurons. A Cre-dependent inhibitory opsin was expressed in SOM interneurons together with GCaMP6f expression in PCs by using combined injection of AAV.DIO.EF1a.eNpHR3.0 and AAV.CaMKII.GCaMP6f into SOM-Cre mice (Fig. 7a). SOM interneurons were silenced during a 5-min period in the middle of the 15-min exploration session, using sequential 10s light pulses (On-bins, λ = 620 nm) separated by 10s intervals (Off-bins, Fig. 7). As shown in the raster plot of Fig. 7b, SOM silencing increased PC spiking, as indicated by the vertical activity bouts synchronized with the onset of the light pulses (Fig. 7b, vertical lines).

**Figure 7.**
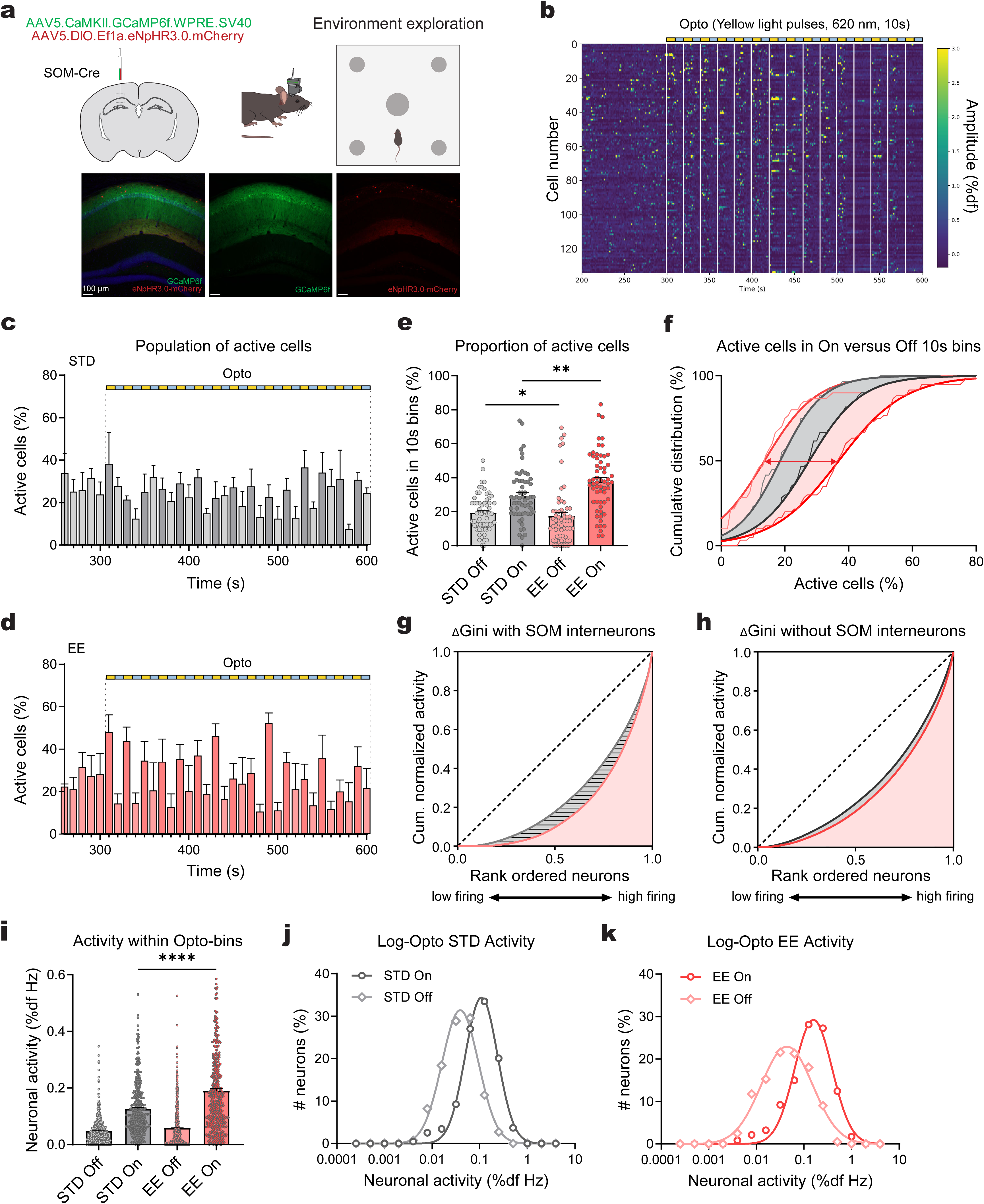
Increased inhibition of CA1 population activity by SOM interneurons. **a)** Schematic illustration showing viral expression of GCaMP6f in CA1 PCs with halorhodopsin (eNpHR3.0-mCherry) expression in SOM cells (left). Confocal images on bottom show GCaMP6f-mCherry labeling (left) and GCaMP6f- (middle) or HR3.0-mCherry-expression (right). **b)** Raster plot of CA1 pyramidal cell responses from an EE mouse. From 300 to 600s, repetitive 10s-light pulses (620 nm LED, On-bins, indicated by the vertical solid lines) were applied, separated by 10s (Off-bins). In total, 15 light stimuli were delivered. Color coded amplitudes range is 0 - 3 %df. **c-d)** Histogram showing the proportion of co-active cells within 10s On- & Off-bins during the Opto-phase in STD (**c**, Off-bins, light grey; On-bins, dark grey, N=4) and EE mice (**d**, Off-bins, light red/pink; On-bins, red, N=4). Optogenetic silencing pulses start at 300s. **e)** Comparison of the average proportion of active cells within 10-s bins during Off and On periods of the opto-block (300-600 s) for STD (Off, light grey; On, dark grey) and EE mice (Off, light red; On, dark red), showing a larger On-bin (p=0.0035, Mann-Whitney test), as well as Off-bin population (p=0.0425, Mann-Whitney test, EE: n=60 bins, N=4 vs STD: n=60 bins, N=4) in EE compared to STD. **f)** Cumulative distribution of the proportion of active cells is small during Off-bins in EE (light red) and STD (light grey) and increases during silencing in On-bins, both for EE (dark red) and STD housing (dark grey, p<0.0001, Wilcoxon matched-pairs signed-rank test). Interestingly, the number of co-active cells during silencing is higher in EE compared to STD (red arrow, p=0.0423, Kolmogorov-Smirnov test, EE: n=60 bins, N=4 vs STD: n=60 bins, N=4). Bold lines represent nonlinear fitting of curves to estimate the shift of the median. **g-h)** Lorenz curves showing more unequal firing across population (Off-bins) in EE (light red) compared to STD mice (light grey, p<0.0001) with unperturbed activity of SOM interneurons (ΔGini with SOM interneurons, **g**). The difference between EE (dark red) and STD (dark grey) was no longer present after SOM silencing (On-bins, ΔGini without SOM interneurons, p=0.4827, **h**). **i)** Comparison of pyramidal cell firing between On- and Off-bins showing higher On-bin activity in EE compared to STD mice (p<0.0001, Mann-Whitney test, EE: n=472 cells, N=4 vs STD: n=436 cells, N=4). **j)** Log-transformed distribution of neuronal activity in STD mice upon SOM silencing is shifted towards higher firing rates without having an effect on the standard deviation of the fitted Gaussian function (F_(1,20)_ = 1.43, p=0.2466). **k)** In EE mice, SOM silencing resulted in both shifting and compression of the log-transformed distribution of neuronal activity towards STD values (F_(1,20)_ = 13.39, p=0.0016). Data are shown as mean ± SEM. * p<0.05, ** p<0.01, and **** p<0.0001.

Analysis of the proportion of co-active cells within 10s bins revealed that the number of active cells increased significantly during the On-bins in both STD and EE animals (Fig. 7c-d). However, the increase in the number of co-active cells during SOM silencing is substantially larger in enriched mice (Fig. 7e). Cumulative distribution of active cells per bin shows that the median proportion of active cells shifts by a factor of 2.7 during SOM silencing in EE animals (from 13.6% to 36.9%, red arrow in Fig. 7f), whereas in STD mice the increase is only 1.5-fold (from 18.5% to 27.7%; n=60 bins, N=4 mice each). This larger shift in the proportion of co-active PCs during SOM silencing indicates that SOM interneurons exert stronger inhibition in enriched mice *in vivo*, consistent with their more efficient recruitment and more powerful inhibitory output described above.

To analyze the effect of SOM interneurons on CA1 PC population dynamics, we calculated the mean firing activity for each neuron separately during Off- and On-bins. A Lorenz curve generated from mean activity during Off-bins, showed a more non-linear distribution in EE compared to STD mice (p<0.0001, n=436 in 4 STD, vs n=472 cells in 4 EE mice), similar to experiments without optogenetics (see Fig. 7g vs Fig. 2f). Also, the Gini index in Off-bins was higher in EE (0.59 vs 0.47). However, during SOM silencing (On-bins), the cumulative distribution of rank-ordered activity no longer differed between EE and STD mice (Fig. 7h, p=0.4827). This indicates that SOM interneurons specifically contribute to the more diverse and more unequal firing distribution of pyramidal cells in enrichment. Furthermore, we quantified the log-normal distribution of firing activity in EE and STD and its shift by optogenetic silencing. Interestingly, silencing of SOM interneurons in STD mice shifted the distribution towards higher firing rates without affecting the standard deviation of the fitted Gaussian function (Fig. 7j, SD_Off_ = 0.275 vs SD_On_ = 0.262, F_(1,20)_ = 1.43, p=0.2466). In EE mice, however, SOM silencing not only shifted firing towards higher rates, but also compressed the standard deviation of the fitted Gaussian distribution towards STD-like values (Fig. 7k, SD_Off_ = 0.387 vs SD_On_ = 0.253, F_(1,20)_ = 13.39, p=0.0016). These findings indicate that the enhanced recruitment of SOM interneurons in EE mice, together with their more widespread lateral inhibitory output observed *in vitro*, plays a critical role in generating more diversity in firing rates and more pronounced sparse coding observed during spatial exploration *in vivo*.

### Improved object-location memory in enriched mice depends on SOM interneurons

To investigate whether the distinct hippocampal information processing observed under enrichment conditions affect learning and memory, we used a novel object location task (OLT, Fig. 8). After habituation to the same arena used for calcium imaging, mice were exposed to two additional new objects and allowed to freely explore them for 15 min. Exploration times during this initial exposure (training) were similar between groups (Supplementary Fig. 7). After 24h, mice were returned to the arena, exposed again to the same objects, with one of them displaced by approximately 30 cm (Fig. 8a). During this test session, both EE and STD mice preferentially explored the displaced object at the novel location (Supplementary Fig. 7). However, the exploration time for the displaced object was longer in enriched mice (68.9% ± 2.5% vs 56.6% ± 1.4%, N=10 and N=9 mice, p=0.0007, Fig. 8b), leading to a 3-times larger discrimination index in EE compared to STD mice (0.38 ± 0.05 vs 0.13 ± 0.03, p=0.0007, Fig. 8c-e).

**Figure 8.**
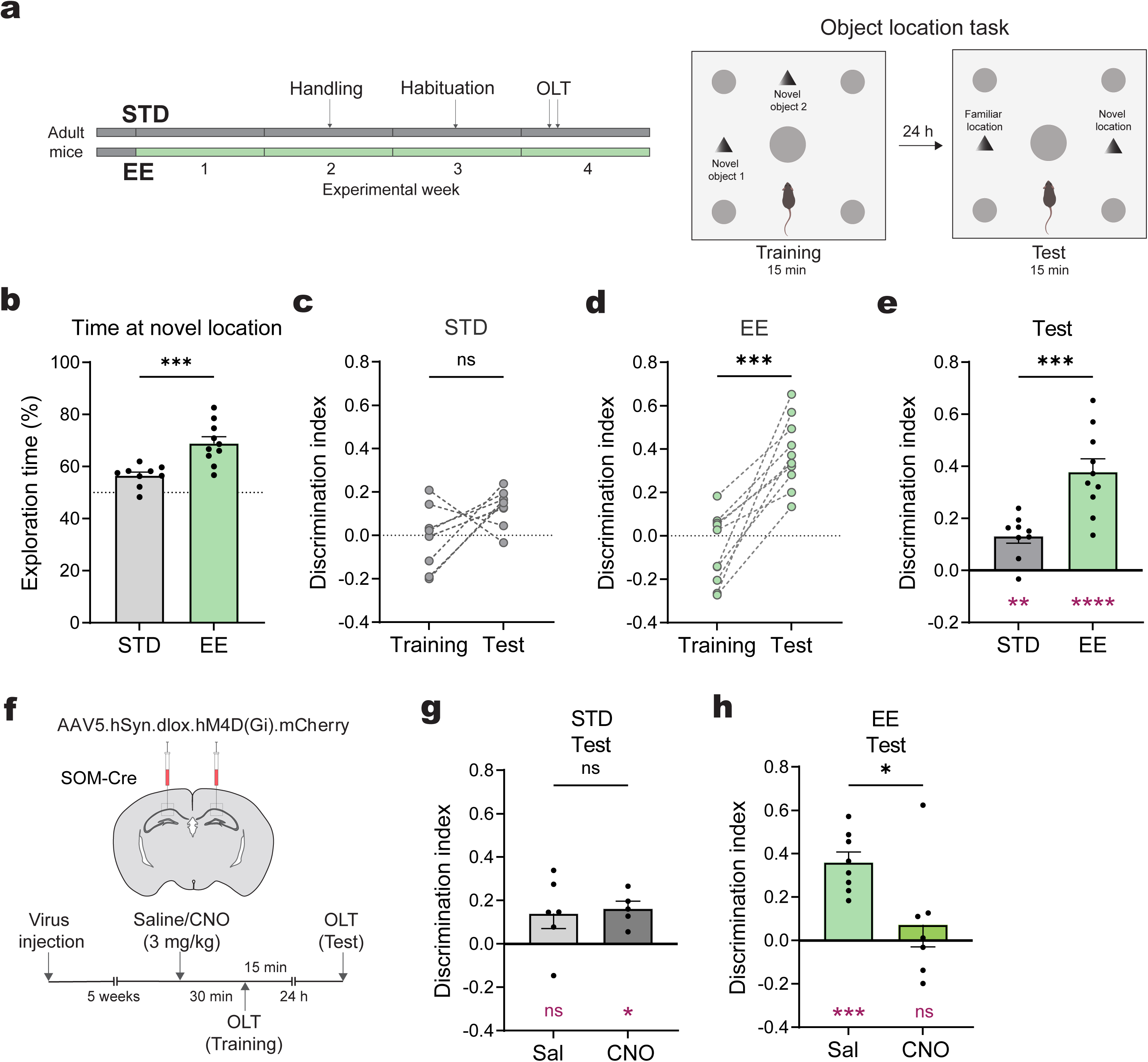
Improved object location memory in enriched mice depends on SOM interneurons. **a)** Left: Experimental timeline. Right: Schematic illustration showing the object location task. During training, mice were exposed to the arena with two novel objects (novel 1 and novel 2). Mice were placed in the arena 24 h later (test) and allowed to explore same objects, with one kept at the same (familiar) location and another displaced to another (novel) location. **b)** Comparison of exploration time of the two objects at familiar and novel locations for STD (grey) and EE (green) mice during OLT (test) shows increased exploration of novel location by EE mice (p=0.0007, unpaired t-test; EE: N=10 vs STD: N=11 mice). **c-d)** Pairwise comparison of discrimination index for STD (**c**) and EE (**d**) during training versus test (p=0.0717, paired t-test, STD-Training vs STD-Test; p=0.0002, paired t-test,EE-Training vs EE-Test). **e)** Discrimination index during OLT (test) is higher for EE (green) compared to STD (grey) mice (p=0.0019, unpaired t-test; same data as in **b-d**). Each group was tested against the theoretical mean (0) using one-sample t-test (shown below in magenta). **f)** Chemogenetic silencing of SOM interneurons in CA1 via virus injection of inhibitory DREADD (hM4D(Gi)-mCherry) into SOM-Cre mice. About 5 weeks after the injection, mice performed OLT. 30 min prior to exposure to the arena with the objects during training, mice were injected either with saline or CNO (3 mg/kg). 24 h later, behavioral testing was performed. **g)** Discrimination index was not affected in STD mice after chemogenetic silencing of SOM interneurons (p=0.7984, unpaired t-test, STD-CNO, N=6 vs STD-Sal, N=5). Each group was tested against the theoretical mean (0) using one-sample t-test (shown below in magenta). **h)** Chemogenetic silencing of SOM interneurons leads to disrupted memory of object location in EE mice (p=0.0200, unpaired t-test, EE-CNO vs EE-Sal, N=7 mice in each group). Each group was tested against the theoretical mean (0) using one-sample t-test (shown below in magenta). Dots represent individual animals. Data are shown as mean ± SEM. ns, not significant, * p<0.05, and *** p<0.001.

To examine the role of SOM interneurons in the improved object location memory, we injected a Cre-dependent inhibitory DREADD virus (AAV.hSyn.dlox.hM4Di-mCherry) into SOM-Cre mice (Fig. 8f). SOM interneurons were then pharmacologically silenced using the DREADD agonist clozapine-N-oxide (CNO) during object learning. Saline-injected controls showed a similarly increased discrimination index in enriched mice compared to STD mice (p=0.0194, Fig. 8g-h). Strikingly, pharmacogenetic silencing of SOM interneurons disrupted object location memory in EE mice and reduced the discrimination index by a factor of 5.1 (EE-CNO: 0.07 ± 0.10 vs EE-Sal: 0.36 ± 0.05, N=7 each, p=0.0200, Fig. 8h). On the other hand, SOM silencing had no detectable effect in STD animals, in which discrimination index remained similarly low under both conditions (STD-CNO: 0.14 ± 0.07, N=6 vs STD-Sal: 0.16 ± 0.03, p=0.7984, N=5, Fig. 8g).

Taken together, these findings show that enrichment experience not only increases the recruitment of SOM interneurons and enhances lateral inhibition between CA1 pyramidal cells, but also promotes diversity in pyramidal cell firing and sparsification of the CA1 population code via SOM interneuron-mediated inhibition. As a consequence, the increased selectivity of pyramidal cell firing in enriched mice allows for a more reliable distinction of different memory items, thereby improving hippocampus-dependent learning and memory.

## Discussion

Our results provide a rigorous multilevel analysis of the consequences of enrichment experience for hippocampal information processing. At the synaptic level, we found not only increased glutamatergic excitation of CA1 pyramidal cells, but also enhanced excitatory synaptic recruitment of SOM interneurons. At the circuit level, we observed increased and more widespread inhibitory output of SOM interneurons as well as stronger and more widespread lateral feedback inhibition between CA1 pyramidal cells. Calcium imaging *in vivo* revealed that population coding in CA1 is sparser and more diverse in enriched mice, showing a larger Gini index and lower number of co-active cells. At the same time, place field firing shows higher spatial selectivity, higher peak firing and increased spatial information. Although optogenetic SOM silencing demonstrates that these interneurons control firing in both STD and enriched mice, SOM-mediated inhibition is substantially stronger under enrichment conditions, where it specifically contributes to the higher Gini index and the broader log-normal distribution of pyramidal cells firing. Finally, using pharmacogenetic manipulation, we show that enhanced SOM-mediated inhibition also enhances hippocampus-dependent discrimination of newly learned memory items.

Environmental enrichment has been extensively studied over the past years, mostly focusing on improved learning, enhanced glutamatergic plasticity and increased synapse density in pyramidal cells^2,9–12^. These studies supported the notion that a larger number of excitatory synapses promotes excitation and improves learning. However, more recent studies analyzing immediate early genes did not report increased neuronal activation. Instead, the proportion of cFos-positive neurons was reduced in enriched mice^13,14^. Because cFos expression is associated with glutamatergic synaptic plasticity^40^, this decrease is surprising. So far, no explanation has been provided for these apparently conflicting observations.

Our study links these apparently conflicting findings, by showing that enrichment experience enhances glutamatergic excitation not only of pyramidal cells but also of GABAergic SOM interneurons. Furthermore, we demonstrate an increased inhibitory output of these cells, suggesting that enrichment enhances synaptic inhibition even more strongly than excitation onto pyramidal cells. This results in an overall lower firing frequency of active pyramidal cells. Importantly, SOM interneurons do not generate a uniform blanket inhibition. Instead, pyramidal cell activity becomes more selective and more diverse, reflected by higher peak firing of place cells, larger inequality of firing across the population, and the finding that only 13% of active cells account for 50% of the overall spiking activity in enriched mice, compared to 20% highly active cells in STD mice. Thus, SOM-mediated feedback inhibition appears to be specifically tuned to support higher spatial information in place cell firing, reduced out-of-field firing, thereby rendering the hippocampal population code more selective and more informative.

Why do SOM interneurons broaden the log-normal distribution of firing frequencies? As suggested previously, a log-normal distribution of firing indicates that the different sources of variability across neurons do not simply sum up, but are instead multiplied through non-linear integration of postsynaptic potentials or nonlinear EPSP-spike transformation^21^. For example, the functional properties of voltage-gated sodium and potassium channels typically generate a log-scale EPSP-spike transformation curve, characteristic for nearly all types of neurons. In addition to these relatively stable intrinsic properties, another more dynamic multiplicative process can arise from shunting inhibition. Because SOM interneurons recruit non-linear α5-subunit-containing GABAA receptors, which can powerfully inhibit dendritic NMDA spikes via shunting inhibition^32,33^, they generate divisive instead of subtractive inhibition. As shunting can be spatially very selective down to the level of single dendritic branches^32^, more widespread feedback inhibition would be expected to increase diversity of pyramidal cell firing and broaden the log-normal distribution of population activity.

What is the functional role of the more pronounced sparse coding observed under enrichment? Sparse coding, characterized by a small group of highly active cells together with a large group of weakly active cells, has been suggested to provide several interesting advantages for neuronal population coding^21–23,43^. First, a small group of about 10% of highly active cells with high spatial information might be sufficient to represent a backbone for an ‘easy’ memory task, generating a ‘good guess’ for matching behavior to many everyday situations^44^. Second, for more demanding and precise pattern discrimination tasks, an increasing number of sparsely active cells could be evaluated, thereby enabling more accurate and precise task performance. Third, if sparse coding supports correct discrimination of related memory items, a mechanism which links sparse coding to experience would be required. Our data provide such a mechanism by showing that experiencing a more diverse environment (larger group size, tunnels, houses), induces the growth of inhibitory synapses to dynamically enhance diversity in neuronal population activity, leading to improved pattern discrimination capabilities. Finally, previous studies have shown that sparsely active pyramidal cells are preferentially recruited into new memory engrams, whereas strongly active cells typically exhibit relatively little remapping^43^. We show that the increasing pool of sparsely active cells in enriched animals is shaped by experience via new excitatory and inhibitory synapses. This growing pool of sparsely active cells and its underlying synaptic connectome might be very useful for binding new information into a unified growing ‘knowledge structure’, representing associated items in overlapping networks^22^. Thereby, our findings provide a mechanism by which hippocampal circuits support the continuous integration of new related memories.

Taken together, our data explain how sparse coding is improved during enrichment experience. In addition to the well-established increase in glutamatergic synapses between principal cells, we have discovered a strong increase in excitatory synapses onto SOM interneurons, accompanied by enhanced and spatially more widespread feedback inhibition between CA1 pyramidal cells. This inhibitory circuit plasticity increases the diversity, selectivity and information content of pyramidal cell firing. As a consequence, less neurons are simultaneously active, thereby improving the overall population sparseness. These findings suggest that diversity in experience shapes the brain to promote diversity in coding. This might be critically important for continuous relational learning and knowledge acquisition in a changing environment throughout life.

## Acknowledgements

We would like to thank Peter Scheiffele for helpful discussions and providing viral constructs for FingR-anti PSD95 intrabodies. We thank Alexandra Lumpi, Leonie Bischofberger and Selma Becherer for mouse genotyping, histochemical staining, confocal imaging and excellent technical assistance. We thank Jan Gründemann for providing help with establishment of miniscope calcium imaging and the Biocenter mechanical workshop for excellent technical support. We would also like to thank the members of Bischofberger lab for helpful discussions. This work was supported by the Swiss National Science Foundation (SNSF, Project 205198 and Project 10006776). The authors declare no competing financial interests.

## Author contributions

E.V., S.K. and J.B. conceived the study and designed the experiments. E.V. performed calcium imaging, learning behavior, cFos and FingR intrabody labeling experiments, immunohistochemistry, confocal imaging and related data analysis. J.I.G. performed patch-clamp analysis of EPSCs in CA1 PCs and mEPSCs in SOMs. S.K. performed all other patch-clamp experiments and analysis. J.B. developed all computational routines and scripts for imaging data analysis. E.V., S.K. and J.B. wrote the manuscript with contributions from all authors.

## Materials and Methods

### Animals

Adult wild-type mice (C57BL/6J) and the following transgenic mice of both genders were used: SOM-IRES-Cre (SST ^tm2.1(cre)Zjh^/J) and SOM-tdTomato (offspring from crossing of heterozygous SOM-Cre with homozygous LoxP-tdTomato (B6.Cg-*Gt(ROSA)26Sor^tm9(CAG-tdTomato)Hze^*/J)) mice. All transgenic mice were obtained from The Jackson Laboratory and were bred by pairing heterozygous transgenic mice with wild-type C57BL/6J animals.

Animals were divided into two groups: 1) for standard housing (STD, 3-4 mice per group), mice were placed in regular individually ventilated cages (IVC, GM500, Tecniplast; 39.1 x 19.9 x 16.0 cm). For enriched environment (EE, 3-6 mice per group), mice were housed in larger rat IVC cages (GR900, Tecniplast; 39.5 x 34.6 x 21.3 cm) with enrichment consisting of 1 running wheel and 3 objects (2 different huts, 1 tunnel), for 2-6 weeks as indicated. Animals were kept in a 12 h light/dark cycle with access to food and water *ad libitum*. Experiments were carried out during the light cycle. All experiments were approved by the Basel Cantonal Committee on Animal Experimentation according to federal and cantonal regulations.

### Stereotaxic virus injections

Mice were injected subcutaneously (s.c.) with carprofen (Rimadyl; Zoetis, 5 mg/kg) as analgesic and anesthetized with isoflurane (4% in O_2_ for induction, 1-2% for maintenance) and placed in a stereotaxic frame (Model 1900, Kopf Instruments). After fixation of the mouse head using ear bars, the skin was cleaned with betadine (Bichsel) and a 1:1 mixture of lidocaine/bupivacaine (Sintetica) was administered s.c. to provide local analgesia. Eye ointment (Bepanthen, Bayer) was applied to eyes to prevent drying. After that, we made an incision to the skin and then levelled the skull relative to bregma and lambda. Next, we made a 0.35-0.4 mm craniotomy above CA1 using different coordinates as outlined below. For injection, we used glass micropipettes (BLAUBRAND Intramark micropipettes) that were pulled using a vertical puller (Narishige). Virus was injected via 5-15-ms-long pulses delivered at 1 Hz using Picospritzer III (Parker Hannifin) and TTL-frequency generator (Voltcraft). After AAV delivery in each injection site, the pipette was held in the brain for at least 5 min before slowly withdrawing. The skin was then sutured using silk thread (Sabana). During surgery, temperature was monitored and adjusted to 36.6 °C using closed-loop heating pad. Analgesia was provided pre- (1 day before surgery) and post-operatively (for 3 days after surgery) in drinking water (carprofen, 5mg/100ml). Post-operatively, health status check and weight measurements were performed.

### Virus injection for *in vitro* electrophysiology and FingR intrabody labeling

For activating GABAergic SOM afferents, SOM-Cre mice were bilaterally injected with ∼100 nl per site of rAAV5.EF1a.DIO-hChR2(H134R).mCherry (titer in vg/ml: 5.2 x 10^12^, UNC vector core, North Carolina, USA). For activating PC axons, C57BL/6J mice were bilaterally injected with ∼100 nl per site of ssAAV5/2.mCaMKIIa.hChR2(H134R).mCherry (v204-5, titer: 5.0 x 10^12^, VVF Zurich, Zurich, Switzerland) at multiple sites (coordinates: AP ±3.3; ML ±3.4; DV -3.75, -3.5, -3.25, -3.0 mm and AP ±3.3; ML ±3.0; DV -3.5, -3.25, -3.0, -2.75 mm relative to bregma). For FingR intrabody labeling experiments, SOM-tdTomato mice were bilaterally injected with 100 nl per site of AAV9.ZnF.hSyn.DIO.PSD-95FingR.mGreenLantern (titer: 2.7 x 10^13^; Peter Scheiffele, Biocenter, Basel) at different sites (coordinates: AP ±2.3; ML ±1.5; DV -1.3, -1.2, -1.1 mm).

### Virus injection for *in vivo* Ca^2+^ imaging and optogenetic and chemogenetic silencing

To express a genetically encoded Ca^2+^-indicator in neuronal populations of CA1 area for *in vivo* Ca^2+^ imaging, C57BL/6J mice were unilaterally injected with 400-500 nl of AAV5.CaMKII.GCaMP6f.WPRE.SV40 (AV7593, titer 4.8 x 10^12^; University of North Carolina (UNC) Vector Core) using the following coordinates: AP -2.3; ML -1.5; DV -1.3, -1.2, -1.1 mm (relative to bregma). For simultaneous imaging and optogenetic silencing experiments, SOM-Cre mice were injected with a mixture of AAV5.CaMKII.GCaMP6f.WPRE.SV40 and AAV5.DIO.Ef1a.eNpHR3.0.mCherry (AV4832D, titer: 4.7x 10^12^; UNC Vector Core) (at a ratio of 1:1; 500 nl in total) using the same coordinates as for imaging. For chemogenetic silencing of SOM interneurons, SOM-Cre mice were bilaterally injected with 400 nl of ssAAV5.hSyn1.dlox.hM4D(Gi).mCherry(rev).dlox.WPRE.hGHp(A) (v84-5, titer 5.6 x 10^12^; VVF Zurich) using the same coordinates.

### Miniscope calcium imaging

#### GRIN lens implantation for Ca^2+^ imaging

One week after virus injection, a GRIN prism lens (1.0 mm in diameter & 4.3 mm in length, Inscopix) was implanted during a second surgery under isoflurane anesthesia and analgesia with buprenorphine (Bupaq; Streuli, 0.1 mg/kg), administered s.c. 30-60 min before surgery. Animals were placed in a stereotaxic frame and lidocaine/bupivacaine was injected as described above. A 1.8 mm diameter craniotomy was made with a trephine drill bit (Fine Science Tools), centered 500 µm lateral to the previous injection site (approx. coordinates: AP -2.3, ML -1.0 mm). Next, a 1.4 mm length posterior-to-anterior cut through the brain tissue 200-300 µm lateral to the injection site and at the depth of 1.9 mm from bregma was made using a cutting blade. The GRIN lens with the prism part oriented in medio-lateral direction was slowly lowered into the brain at the cutting depth of 1.9 mm and fixed to the skull with UV light-curable glue (Loctite 4305, Henkel). Then the skull was sealed with blue light-curable glue (Scotchbond, 3M), wound-closing material (Vetbond, 3M) and dental acrylic (Paladur, Kulzer), and a custom-made titanium head bar (Biocenter mechanical workshop) was attached. To prevent damage of the dorsal surface of the lens overtoping the skull, the lens was covered with silicon. Post-operative analgesia was applied for 3 days, and the health status check was carried out daily for a week. Mice were allowed to recover for about 10 days before checking for fluorescent signal and viral expression level.

#### Microscope baseplate fixation for miniature microscope imaging

Before miniature microscope imaging experiments, a microscope baseplate (Inscopix) was attached to the head of a mouse. Briefly, mice were head-fixed at the head bar on a flying saucer style running wheel in a custom-built mounting station (Thorlabs). The base plate was attached to the microscope (nVoke 2.0, Inscopix) and the microscope’s objective was positioned above the lens surface. The position was then adjusted accordingly to find a field of view (FOV) with optimal fluorescent signal from cells. Mice were briefly anesthetized with isoflurane and the baseplate was glued to the dental cement using blue light-curable glue (Vertise Flow) and the lens was protected using a baseplate cover (Inscopix).

#### Handling and habituation

After the baseplate fixation, mice were handled for a few days before imaging experiments. In addition, each day mice were habituated to the procedure of mounting the microscope. One day before the experiment, mice were placed into their home cage and a short trial recording was obtained to adjust the FOV for consecutive imaging. Furthermore, the mice were habituated to the open field (OF) recording arena (80 x 80 x 20 cm, PVC), equipped with 5 fixed objects, for at least one week for 15 min per day. Spatial cues were fixed on the inner side of one wall and on three locations surrounding the arena.

#### Ca^2+^ imaging and optogenetics

For simultaneous imaging and optogenetic silencing experiments, a nVoke system (nVoke 2.0, Inscopix) was used together with a nVision camera system (Inscopix) to record mouse behavior. To ensure free movement of animals in an environment, we used a cable commutator (Inscopix) which was centered at a height of about 1m above the mouse. To record Ca^2+^ activity, the Inscopix Data Acquisition Software (IDAS) was used. Ca^2+^ fluorescence was imaged at a frame rate of 20 Hz with a blue LED power (EX-LED, λ_exc_ = 455 ± 8 nm) set to 0.4–0.9 mW/mm^2^ and the detector gain to 1.0–4.0. The gain was adjusted for each animal according to the expression level. For optogenetic silencing experiments, a red LED was used (OG-LED, λ_exc_ = 620 ± 30 nm) with the power level set to 80% (4 mW/mm^2^). The FOV size was 1280 x 800 pixels corresponding to 1050 x 650 µm.

Before each session, mice were placed on a running wheel in a small custom-built setup and head-fixed at the head bar. The microscope was attached to the animal’s head via the baseplate. Mice were then placed into the familiar recording arena (80cm x 80 cm) for a duration of 15 min per session. For optogenetic silencing of SOM interneurons, the session was divided into 3 continuous time blocks lasting 5 min each. Optogenetic pulses were applied only in the middle of the 3 blocks (pre, opto, post), while recording of Ca^2+^ activity was performed continuously for the full 15 min session. During the opto block, 15 light stimuli with a duration of 10 s (On-Bin) were delivered at the interval of 10 s (Off-Bin) simultaneously with recording.

#### Imaging data analysis

Miniscope imaging data were analyzed using Inscopix Data Processing Software (IDPS, Inscopix) and custom-written scripts in Python. First, data from all sessions were combined in single time series and preprocessed in IDPS. Preprocessing included cropping, temporal and spatial down sampling by a factor of 2, spatial bandpass filtering and motion correction (relative to a reference frame). For further processing, sessions were separated and dF/F videos were generated. Cell detection was performed using principal and independent component analysis (PCA-ICA). For this analysis, we first estimated the number of individual components (ICs) from a maximum dF/F_0_ intensity projection and initialized the PCA-ICA algorithm with the number of principal components (PCs) set to 120% of ICs for automatic extraction of z-scored fluorescence traces. These automatically detected cells were visually inspected and units with diameters >70 µm were excluded from the cell set.

The detected cell set was then used to calculate regions of interests (ROIs) for extraction of relative change in somatic fluorescent signals of individual neurons from the motion-corrected videos. Specifically, ROIs were defined by drawing a contour line at 50-60% of the peak fluorescence intensity in the center of the soma. Furthermore, we rejected cells in case the area overlap with a neighboring cell was more than 20%. The baseline fluorescence (F_0_) of each cell was calculated from the median of the time-dependent fluorescent trace, which was possible because hippocampal pyramidal neurons show sparse firing activity with sufficient periods of resting calcium concentration (see Figure 1). Finally, the relative change in Ca^2+^-concentration was quantified as (f(t)-F_0_)/F_0_ (%df), by subtracting baseline fluorescence F_0_ from time-dependent fluorescence signal f(t) and normalizing it to F_0_. For detection of Ca^2+^ events, traces were low pass filtered using a mild Savitzky-Golay filter to preserve peak amplitudes. Peaks were accepted if they were larger than 2-times the standard deviation of the whole trace and larger than a minimal threshold of 0.8 %df, to avoid false positive events due to noise.

#### Analysis of neuronal activity

Event frequency was calculated by dividing the number of events by recording time. The mean amplitude per cell was estimated by averaging the amplitudes of all events and average spiking activity was calculated for each cell as the product of event amplitude and event frequency. For temporal population analysis, time was separated into 10-s time bins (in total, 90 bins for 900 s) and the number of active cells during such a bin was divided by the number of all cells which were active at least once. For the analysis of optogenetic experiments, On- and Off-bins between opto-start and opto-stop (300s-600s) were analyzed separately. To analyze the log-normal distribution of spiking activity and event amplitude, data were transformed to logarithmic scale. The binned distribution was the fitted with a gaussian function in GraphPad Prism. To generate the Lorenz curve, neurons were first rank ordered according to their activity, and the cumulative sum was calculated as a function of neuron number, both normalized to one. Finally, to calculate the Gini index (G), we first numerically calculated the area under the Lorenz curve A, subtracted this from the area below the identity line (0.5), and normalized the value to range between 0-1: G = (0.5 – A) / 0.5.

#### Place field firing

To analyze place field firing in CA1 pyramidal cells, we first analyzed the track of a mouse within the arena (80cm x 80cm) using the nose point obtained using EthoVision XT software (Noldus, version 17.5). Second, we calculated the instantaneous velocity and applied a 1s temporal box-car low-pass filter. This allowed to exclude resting activity from exploration with velocities lower than 2cm/s. Afterwards, calcium activity was binned in 2D for each cell using a bin size of 1.2cm x 1.2cm, followed by spatial low-pass filtering using a scipy gaussian filter with a standard deviation of 2 bins. Place field borders were calculated using a contour line at 30% of the maximal calcium signal in the center of the place field. Firing fields were accepted as place fields in case the peak calcium amplitude was larger than 2-times the standard deviation across the whole arena and larger than a minimal threshold of 0.2%df. Furthermore, the area was required to be larger than 78cm^2^ corresponding to a minimum circular diameter of 10 cm. Spatial sparsity was calculated according to Skaggs et al. (1996) using the spatial inhomogeneity of the activity ***r_i_*** at bin ***i*** with occupancy ***p_i_*** as:

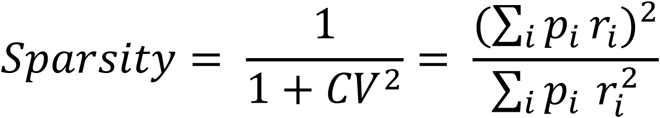

Furthermore, the Shannon information **SI** was calculated according to Skaggs et al. (1996) as

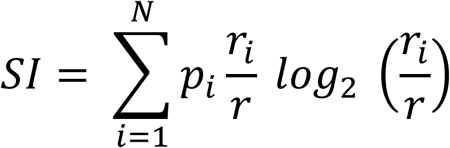

with the activity ***r_i_*** at bin ***i*** with occupancy ***p_i_***.

### Novel object location task

To evaluate hippocampus-dependent spatial memory in mice, we performed novel object location task (OLT). This task is based on spontaneous tendency of mice to explore objects at novel rather than familiar locations. Prior to OLT, mice from STD and EE groups were handled by experimenter daily for 5-10 min for 3-5 days. For EE mice, handling begun about 1 week after the start of the EE housing. Handling was followed by habituation to the OF arena (80 x 80 cm). Mice were transferred to the room about 30 min before the beginning of an experiment for acclimatization.

The task consisted of two phases: training/sample phase and test phase. During the training phase, two identical objects were placed into the OF arena at a fixed location of 15 cm from the arena’s wall. The objects were custom-made white or black pyramids (4 x 4 x 6 cm, polyoxymethylen)^45^. Spatial cues were fixed on the inner side of one wall and on three locations surrounding the arena. Mice were placed into the arena and allowed to explore the objects for 15 min. During the test phase (24 h later), one of the objects was displaced by 34 cm to a novel location (15 cm from the wall). Mice were allowed to explore the objects for 15 min. The arena and the objects were cleaned thoroughly with 70% ethanol between animals to remove olfactory cues. The behavior was recorded using a CMOS-camera (nVision, Inscopix) centered above the arena at a height of about 1.2 m. The recording was done at 30 frames per second and 1920 x 1080 resolution. The analysis of the behavioural videos was performed using EthoVision XT. The exploration time of objects at different locations was defined by tracking the nose of mouse directed or sniffing an object at a distance of no more than 2 cm.

The Discrimination Index DI was calculated for training and test phases using the following formula:

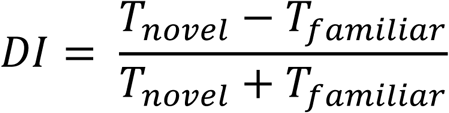

Because both objects were novel at the training session, the DI training should be close to zero. Animals were excluded from the analysis (1 out of 20 mice) if they showed an initial bias toward one object, as indicated by a DI _Training_ of less than -0.3 or more than 0.3, respectively.

### Chemogenetic manipulation of neuronal activity during OLT

About 3-4 weeks after the virus injections of AAV5.hSyn1.dlox.hM4D(Gi).mCherry, mice were handled for 3-5 days and habituated to the OF arena for about a week. To specifically inhibit SOM cells during the training/sample phase, mice were injected i.p. on day 1 of OLT either with saline (STD-Sal; EE-Sal) or with 3 mg/kg of a water-soluble version of clozapine-N-oxide (clozapine-N-oxide dihydrochloride, CNO; Tocris, 6329) 30 min before the exposure to the arena with the objects. The test phase after 24 h, was identical to control without CNO.

### Spatial exploration for cFos counting

For cFos labeling, mice from standard housing (STD) and enrichment (EE) were randomly assigned to a group: home cage (HC-STD, HC-EE) and spatial exploration (NE-STD, NE-EE) groups. Mice from a NE group were placed into a Novel Context (Eurostandard type IV cage (59.5 x 38.0 x 20.0 cm), with randomly located objects (3 familiar and 5 novel) and a running wheel) and allowed to explore the environment for ∼1 h. 90 min after the start of exploration, all mice were perfused. Mice from the HC group did not perform any exploration and were taken from their home cage (HC).

### Immunohistochemistry

Mice were anesthetized using ketamine (130 mg/kg) and xylazine (15 mg/kg) administered i.p. and transcardially perfused with approx. 25 ml phosphate buffered saline (PBS) followed by 25-45 ml 4% paraformaldehyde (PFA) in 0.1M PBS. Brains were removed and fixed overnight in 4% PFA at 4°C. After fixation, 100-µm-thick horizontal dorsal hippocampal (for cFos immunostaining) were cut using a vibratome. After washing three times with PBS and blocking for 1 h at room temperature in blocking solution (5% normal donkey serum (NDS) with 0.3% Triton X-100 in PBS), slices were incubated for 12 h at room temperature, followed by incubation for 36 h at 4°C with primary antibodies (for C57BL/6J: rabbit anti-cFos, 226003, Synaptic Systems, 1:1000; rat anti-somatostatin, MAB354, Millipore, 1:250; for SOM-tdTomato: rabbit anti-cFos, 1:1000) in carrier solution (including 5% NDS with 0.3% Triton X-100 in PBS). After washing three times with PBS, secondary antibodies (for C57BL/6J: donkey anti-rabbit Alexa Fluor 488, A21206, Invitrogen, 1:500; donkey anti-rat Alexa Fluor 594, A21209, Invitrogen, 1:500; for SOM-tdTomato: donkey anti-rabbit Alexa Fluor 488, 1:500) with DAPI (1:10000) in carrier solution were applied for 24 h at 4°C. After subsequent washing, slices were embedded in Prolong Gold and mounted on glass slides.

### Confocal imaging for cFos cell counting

Image stacks from CA1 regions from 6 slices (from both hemispheres, resulting in 12 stacks per area per mouse) were acquired with a 10x air objective (Plan-Apochromat, NA 0.45, Zeiss) using a confocal laser scanning microscope (LSM900, Zeiss). Images with the size of 2048x2048 pixels were scanned with Z-step of 4 µm. For cFos counting in SOM+ cells, image stacks were acquired using a 20x air objective (Plan-Apochromat, NA 0.8, Zeiss) at 2048x2048 pixel resolution and Z-step of 1 µm.

Image stacks were analyzed using ZEN 2 software (Zeiss) and ImageJ (Fiji). Prior to analysis, orthogonal maximum intensity projection from the confocal image stacks was obtained for each region using ZEN 2. After background subtraction, cells were counted, using ImageJ macros utilizing CSBDeep and StarDist2D plugins. Cells were counted in a selected area of the projected stack. Visual inspection was done to remove falsely identified objects. Cells with an area less than 45 µm were excluded from the analysis. The number of cFos+ and cFos+SOM+ cells was counted, and then normalized by the total number of CA1 PCs in defined area using DAPI+ signal (Deng et al., 2013; Lamothe-Molina et al., 2022). The number of cFos+SOM+ cells was defined by the presence of green (Alexa Fluor 488) and red (tdTomato) signals, which was then normalized by the total number of SOM+ cells (tdTomato).

### FingR intrabody experiments and synapse counting

Mice injected with FingR intrabody were anesthetized and transcradially perfused with 25 ml PBS followed by 40 ml 4% PFA and 16% Picric Acid in 0.1M PBS. Brains were fixed overnight in 4% PFA with 16% Picric Acid at 4°C. After that, brains were washed 3-5 times in PBS and 120-µm-thick horizontal slices were cut on vibratome. The slices were then incubated with CUBIC-L solution (N-butyl-diethanolamine in water) to remove lipids for tissue clearing for 6 h (the solution was changed once). After washing three times in PBS and then blocking for 1h at room temperature in blocking solution, slices were finally incubated for 96h at 4°C with primary antibody (rabbit anti-vGlut1, Millipore, 1:2000) in carrier solution. Following that, we added a secondary antibody (donkey anti-rabbit Alexa Fluor 647, 1:500) to carrier solution and incubated the slices for 24h at 4°C.

We obtained confocal image stacks of selected SOM interneurons in CA1 oriens/alveus region using a 40x oil objective (Plan-Apochromat, NA 1.3) and the confocal laser scanning microscope. Cells were selected based on bipolar morphology and expression of PSD95-mGreenLantern (EGFP) and SOM-tdTomato (mGreenLantern+tdTomato+). The image size was set to 1024x1024 pixels, Z-step was 0.2 µm. Images were deconvolved and orthogonal maximum intensity projection was acquired using ZEN 2. Subsequent analysis was done using custom-written macro in ImageJ utilizing CSBDeep and StarDist2D. After background subtraction in EGFP channel, synapses in selected dendritic branches of tdTomato+ cell were counted. In addition to the number of synapses, total dendritic length, synapse density and area per synapse were analyzed.

### Electrophysiology

#### Slice preparation for patch-clamp recordings

Slice electrophysiology experiments were performed in mice aged between 6 and 12 weeks. Mice were first anesthetized with isoflurane (4% in O2, Vapor, Draeger) and then immediately decapitated. To increase cell viability, mice were exposed to oxygen-enriched atmosphere for at least 10 min before decapitation. Slices were cut as previously described^46,47^. Briefly, the brain was dissected in ice-cold sucrose-based solution. Horizontal hippocampal brain slices (350 μm thick) were cut at an angle of 20° to the dorsal surface of the brain along the dorsoventral axes of the hippocampus using a Leica VT1200 vibratome. For cutting and storage, a sucrose-based solution was used, containing (in mM): 87 NaCl, 25 NaHCO_3_, 2.5 KCl, 1.25 NaH_2_PO_4_, 75 sucrose, 0.5 CaCl_2_, 7 MgCl2, and 10 glucose (equilibrated with 95% O_2_ / 5% CO_2_, Osmolarity 340 mOsm). Slices were kept in a holding chamber at 35 °C for 30 min after slicing and subsequently stored at room temperature until experiments were performed.

#### Whole-cell recordings

Electrophysiological recordings of acute brain slices were conducted within about 6 hours after slice preparation. Slices were placed into a bath chamber, with continuous perfusion of oxygenated artificial cerebrospinal fluid (ACSF) at room temperature (∼ 25 °C) which contained (in mM): 125 NaCl, 25 NaHCO_3_, 25 glucose, 2.5 KCl, 1.25 NaH_2_PO_4_, 2 CaCl_2_, and 1 MgCl_2_ (equilibrated with 95% O_2_/5% CO_2_, Osmolarity 320 mOsm). CA1 pyramidal cells (CA1 PCs) were visually identified in the pyramidal-cell layer close to the border of striatum radiatum using infrared differential interference contrast (IR-DIC) video microscopy and a 40x objective. CA1 SOM interneurons (CA1 SOMs) were identified using three basic criteria including: location in alveus, bipolar shape and red fluorescence (tdTomato) using a cooled CCD camera system (SensiCam, TILL Photonics, Gräfelfing, Germany). TdTomato-positive SOM interneurons were detected using a light source with an excitation wavelength of 555nm (Polychrome V, TILL Photonics), which was connected to the microscope using a quartz fiber optic light guide. The illumination intensity was kept low to avoid possible photo-bleaching of the neurons and subsequent phototoxicity.

Voltage recordings of CA1 PCs and SOMs were conducted with patch pipettes (3-6 MΩ) pulled from borosilicate glass tubes with 2.0 mm outer diameter and 0.5 mm wall thickness (Hilgenberg, Malsfeld, Germany; Flaming-Brown P-97 puller, Sutter Instruments, Novato, USA). For recordings of the electrically evoked AMPA excitatory postsynaptic currents (EPSCs), the spontaneous or miniature excitatory postsynaptic currents (sEPSCs or mEPSCs) as well as the optogenetically evoked inhibitory postsynaptic currents (IPSCs), the patch pipettes were filled with a physiological Cs-based intracellular solution containing (in mM): 136 Cs-Gluconate, 4 CsCl, 10 EGTA, 10 Hepes, 2 MgCl_2_, 2 Na_2_ATP, 1 Phosphocreatine and 5 QX-314 adjusted to pH 7.3 with CsOH.

Voltage clamp recordings were obtained using a Multiclamp 700B amplifier (Molecular Devices, Palo Alto, CA, USA), filtered at 10 kHz (or 1 kHz for sEPSCs and mEPSC recordings), and digitized at 20kHz with a CED Power 1401 interface (Cambridge Electronic Design, Cambridge, UK). In all voltage-clamp recordings, series resistance (< 20 MΩ) was monitored online and experiments were discarded if Rs changed more than 20%. To examine voltage dependence and time course of postsynaptic currents, whole-cell series-resistance (8–12MΩ) compensation was used (70–80% correction) except for sEPSC and mEPSC recordings. Data acquisition was controlled using IGOR Pro 6.31 (WaveMetrics, Lake Oswego, Oregon) and the CFS library support from CED (Cambridge Electronic Design, Cambridge, UK).

#### Extracellular synaptic stimulation

To electrically stimulate synaptic inputs, stimulation pipettes were filled with a HEPES-buffered sodium rich solution to apply short negative current pulses (10-100 μA, 200 μs). To stimulate Schaffer collaterals (SCs), the pipettes were placed into stratum radiatum (s.r) close to the CA2 - CA1 border at a distance of ≥ 500 μm (500–800 μm) from the recorded neuron. To stimulate CA1 afferents the pipettes were placed into stratum oriens (s.o) ∼ 200 μm away from the recorded SOM interneuron to avoid direct stimulation of its dendritic arbor. Stimulus artefacts have been truncated in most figures for clarity.

#### Channelrhodopsin-assisted inhibitory Postsynaptic Currents (ChR2-IPSCs)

in the postsynaptic PCs were evoked as previously described^32,48^. A diode laser (DL- 473, Rapp Optoelectronic) was coupled to the epifluorescent port of the microscope (Zeiss Examiner, equipped with a 40x objective; Carl Zeiss Microscopy GmbH, Jena, Germany) via fiber optics. The laser was controlled via TTL pulses. For recording SOM-mediated IPSCs onto CA1 PCs, the illumination area was focused on the middle of stratum lacunosum moleculare to optogenetically activate SOM axons. For recording feedback IPSCs onto CA1 PCs, the illumination area was focused on stratum pyramidale to optogenetically activate PC axons. For the i-o curves we used laser intensities of 1–5 mW for 5 ms (increments of 1 mW) with inter-trial interval of 15 sec. For the lateral distribution optogenetic activation of SOM and PC axons, we used a 5 msec single pulse of either 4 mW or 3 mW, respectively, with inter-trial interval of 10 sec. ChR2-IPSCs were recorded in voltage-clamp mode at 0 mV. SOM-IPSCs were recorded in the presence of 10 μM NBQX and 25 μM D-AP5.

#### Pharmacology

Stock solutions of NBQX (Tocris Bioscience, 10 mM), D-AP5 (Tocris Bioscience, 25 mM), Gabazine (SR 95531 hydrobromide, Sigma-Aldrich, 10 mM) and Tetrodotoxin (TTX, Alomone Labs, 1 mM) were dissolved in water. All drugs were stored as aliquots at -20 °C and diluted in ACSF.

#### Patch-Clamp Data Analysis

Patch-clamp data was analysed offline using the open-source analysis software Stimfit (https://neurodroid.github.io/stimfit) and customized scripts written in Python. For analysis of amplitudes, decay time course and paired-pulse ratios of evoked PSCs average traces of at least 5 repetitions were used. The coefficient of variation (1/CV^2^) was defined from CV = SD / Mean of 50 consecutive EPSCs. For calculations of the integral, traces were low-pass filtered (third-order Butterworth, 100 Hz cutoff) to determine the start and endpoint of the integral as the intersection of the smoothed trace with the baseline. The integral was then calculated as the sum of all values of the original trace minus the baseline in between start and endpoint. For the analysis of sEPSCs and mEPSCs a template-matching algorithm, implemented in Stimfit, was used as described previously^49^. Automatically detected events were visually controlled and false positive events were deleted. The remaining events were fitted with the sum of two exponential functions revealing the amplitude, rise time and decay time of the events.

### Statistics

Statistical analysis was performed using GraphPad Prism version 11 (GraphPad Software, La Jolla, CA). No statistical methods were used to pre-determine sample sizes. Data distributions were assessed for normality using Shapiro-Wilk test.

For comparison between two independent groups, two-tailed unpaired Student’s t-tests were used when data were normally distributed. When paired measurements were obtained from same cells or animals, two-tailed paired t-tests were applied. For non-normally distributed data, the non-parametric Mann-Whitney U test or Wilcoxon matched-pairs signed-rank test was used for comparisons of unpaired or paired data sets, respectively.

For comparisons involving multiple groups or factors, analysis of variance (ANOVA) followed by appropriate post hoc tests was used, as specified in figure legends or main text. Differences between cumulative distributions were assessed using a Kruskal-Wallis test. Kolmogorov-Smirnov test was used to compare distributions of the synaptic parameters such as rise time and latency.

For analyses of Lorenz curves and firing rate distributions, cumulative activity distributions were compared using a Kolmogorov-Smirnov test. Inequality of firing across neurons was quantified using the Gini index. Log-normal firing distributions were analysed after logarithmic transformation of firing activity. Gaussian functions were fitted to raw or log-transformed data and differences between fitted distributions were evaluated using a sum-of-squares F-test.

Group data are presented as mean ± SEM, unless otherwise stated. Statistical values are reported in the main text or figure legends. Significance levels are denoted as *P<0.05, **P<0.01, ***P<0.001, ****P<0.0001.

## Supplementary Figure Legends

**Supplementary Figure 1:**
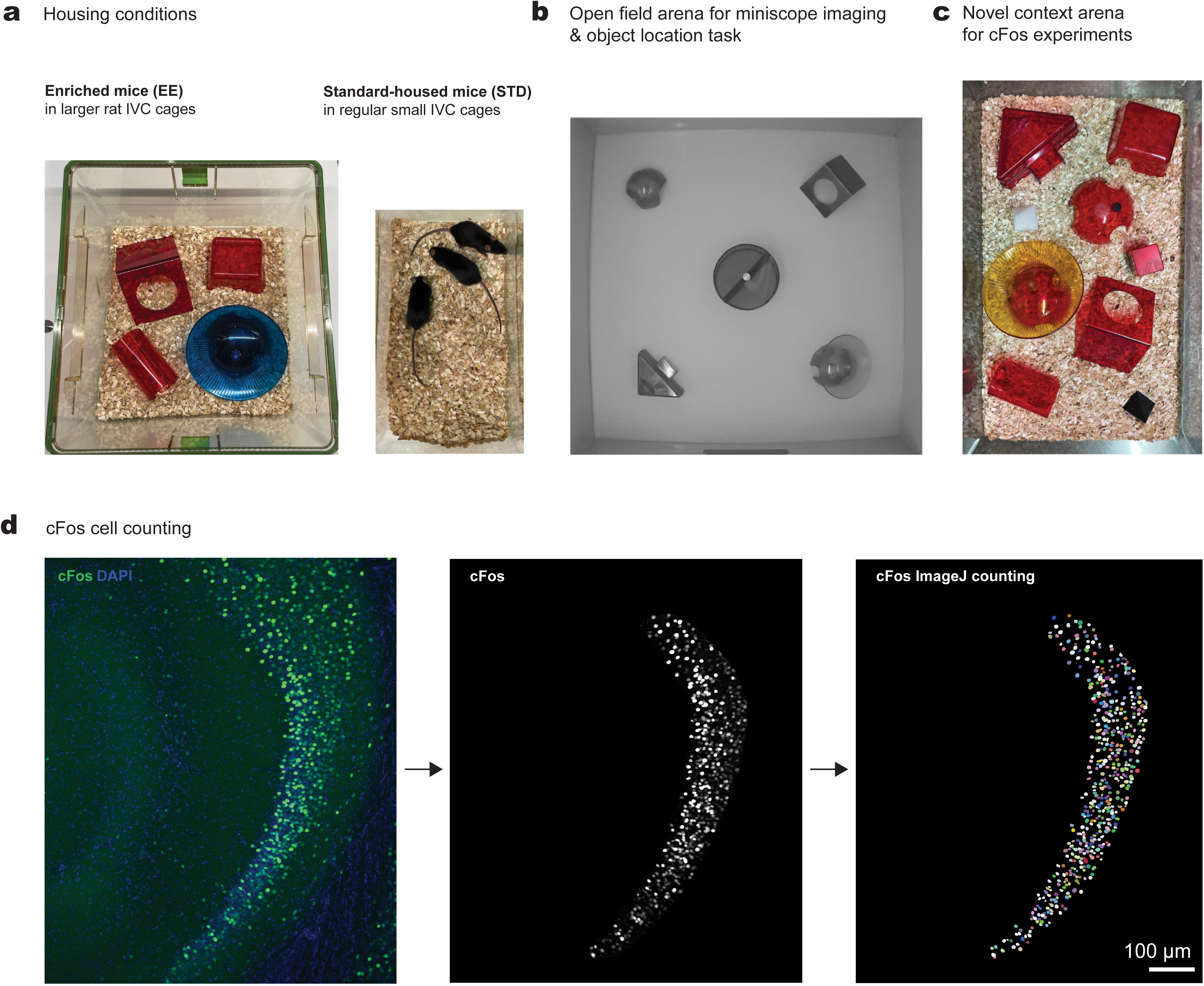
Housing conditions and recording arenas used for imaging and cFos experiments. **a)** Example images of the housing conditions for STD (left) and EE (right) mice **b)** Schematic of recording open field arena used for *in vivo* miniscope imaging experiments. **c)** Example image of a novel context arena used for cFos labeling experiments. **d)** Example images showing cFos cell counting procedure in ImageJ: area selection, background subtraction, cFos cell identification and counting.

**Supplementary Figure 2.**
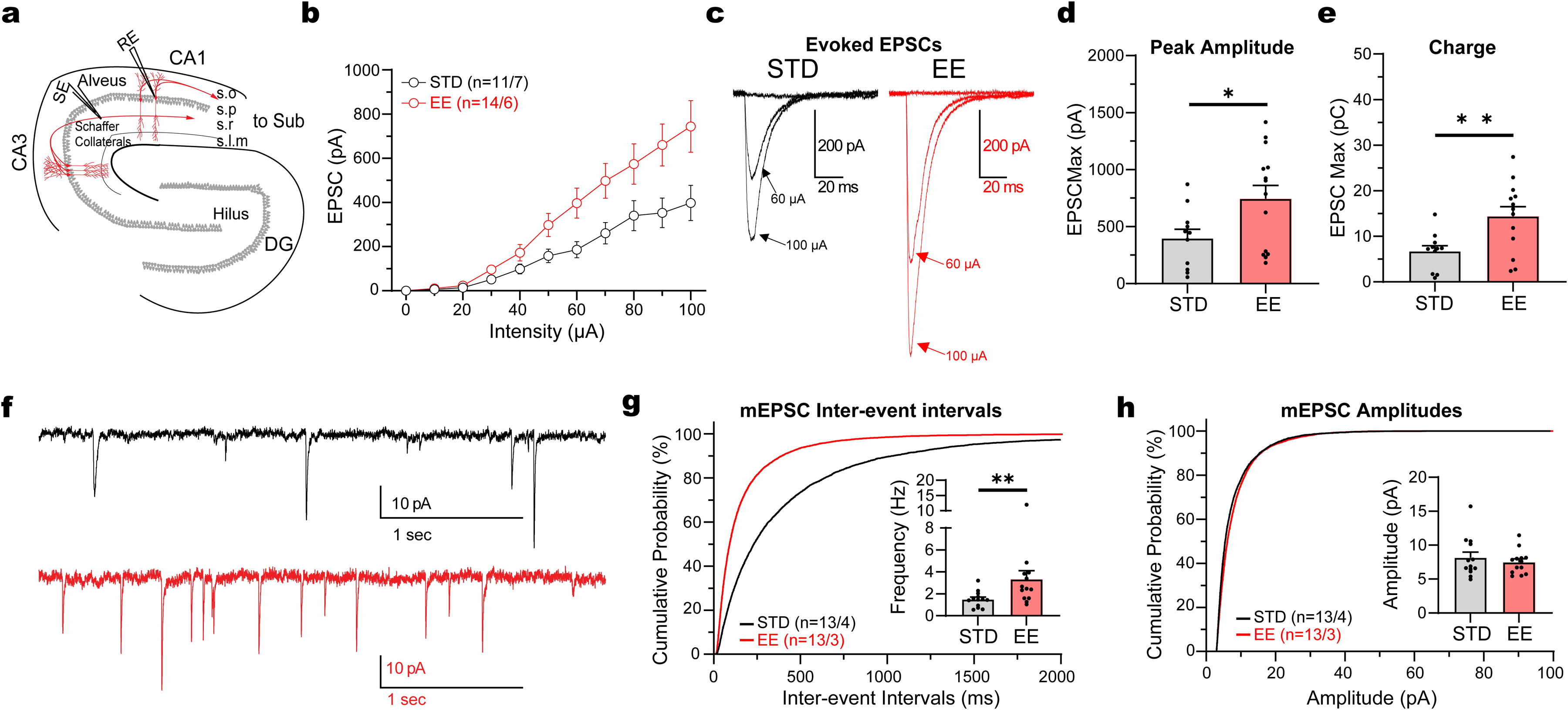
Enrichment increases excitatory drive onto CA1 PCs. **a)** Schematic illustration showing electrical stimulation of Schaffer collaterals while recording evoked EPSCs in CA1 PC. **b)** Stimulus-response curve showing larger amplitude of evoked EPSCs in response to increasing electrical stimulation in Schaffer collaterals in CA1 PCs from EE mice (n=14 cells, N=6 mice) as compared to STD housing (n=11 cells, N=7 mice) in presence of gabazine (p<0.0001, 2-way ANOVA, F _(23, 230)_ = 12.98). Stimulation intensities were ranging from 0 to 100 μA. **c)** Representative traces of evoked EPSCs in CA1 PCs at different intensities of 0, 60 and 100 μA from a STD (black) and an EE mouse (red). **d)** Larger evoked maximal EPSC amplitudes in CA1 PCs in EE compared to STD housing (p=0.0301, Unpaired t-Test). **e)** Larger evoked EPSC integrals in CA1 PCs in EE compared to STD housing (p=0.0077, Unpaired t-Test). **f)** Representative traces of miniature EPSCs (mEPSCs) from STD (black) and EE (red) mice recorded at -70 mV in presence of gabazine and TTX. **g)** Cumulative frequency distributions of interevent intervals of mEPSCs recorded from STD (black) and EE (red) CA1 PCs (P<0.0001, Kruskal-Wallis test, EE: n=13 vs STD: n=13). Inset: Increased mEPSC frequencies per cell in EE mice (p=0.0042, Mann-Whitney test). **h)** Frequency distribution of amplitudes of mEPSCs recorded from STD (black) and EE (red) CA1 PCs. Inset: average mEPSC amplitudes per cell (STD: 8.15 ± 0.82 pA vs EE: 7.48 ± 0.49 pA, p=0.9598, Mann-Whitney test). Data are presented as mean ± SEM. * p<0.05 and ** p<0.01.

**Supplementary Figure 3:**
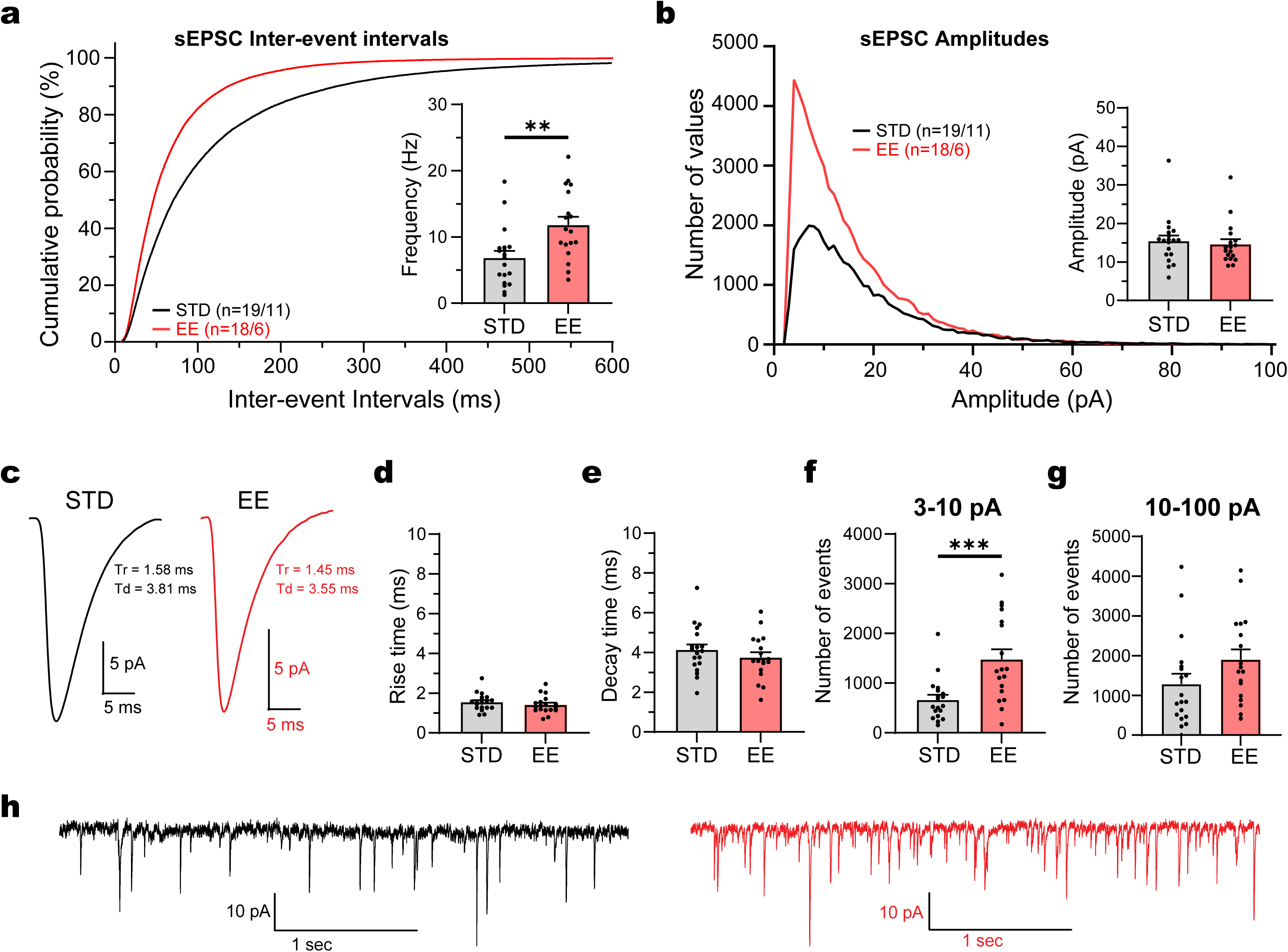
Enrichment enhances the frequency of sEPSCs in CA1 SOMs. **a)** Cumulative frequency distributions of interevent intervals of spontaneous EPSCs (sEPSCs) recorded from STD (black) and EE (red) SOM cells (P<0.0001, Kruskal-Wallis test). Inset: average sEPSC frequencies per cell (STD: 6.86 ± 1.02 Hz vs EE: 11.82 ± 1.23 Hz, p=0.0036, Unpaired t-Test). **b)** Frequency distribution of amplitudes of sEPSCs recorded from STD (black) and EE (red) SOM cells. Inset: average sEPSC amplitudes per SOM cell (STD: 15.45 ± 1.45 pA vs EE: 14.62 ± 1.31 pA, p=0.3275, Mann-Whitney test). **c&h**) Representative average sEPSC trace (c) and (h) Representative traces of sEPSCs from STD (black) and EE (red) mice recorded at -70 mV in presence of gabazine. **d)** Average sEPSC rise times per SOM cell (STD: 1.55 ± 0.10 ms vs EE: 1.42 ± 0.10 ms, p=0.3467, Unpaired t-Test). **e)** Average sEPSC decay times per SOM cell (STD: 4.13 ± 0.27 ms vs EE: 3.74 ± 0.26 ms, p=0.3133, Unpaired t-Test) **f&g)** Average number of events (F) for 3-10 pA amplitudes (STD: 664.9 ± 98.5 events vs EE: 1479.0 ± 203.3 events, p=0.0008, Mann-Whitney test) and (G) for 10-100 pA amplitudes (STD: 1286.0 ± 257.9 events vs EE: 1899.0 ± 256.6 events, p=0.0569, Mann-Whitney test) per SOM cell. Data are presented as Mean ± SEM. ** p<0.01 and *** p<0.001. N=11 animals, n=19 SOM cells in STD vs N=6 animals, n=18 SOM cells in EE.

**Supplementary Figure 4:**
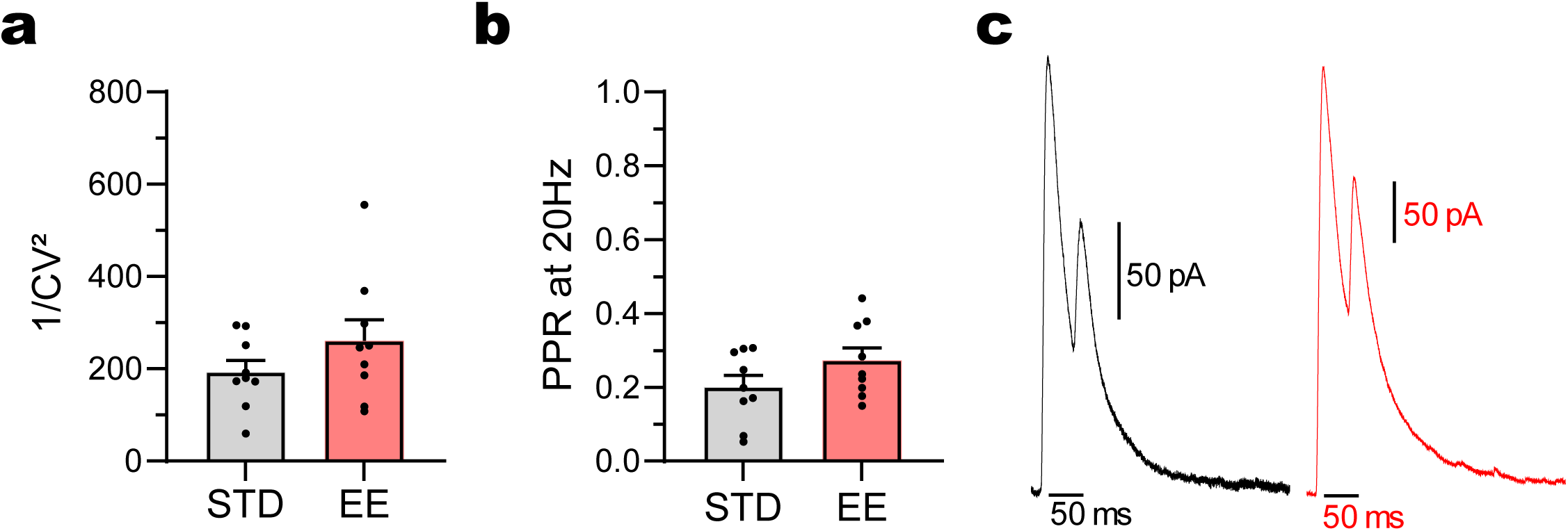
PPR and CV of SOM-mediated IPSCs are not affected by enrichment. **a&b**) Quantification of (A) 1/CV² (STD: 192.7 ± 25.8, N=6 animals, n=9 PCs vs EE: 260.3 ± 45.9, N=4 animals, n=9 PCs, p=0.3401, Mann-Whitney test) and (B) PPR (STD: 0.20 ± 0.03, N=6 animals, n=9 PCs vs EE: 0.27 ± 0.03, N=4 animals, n=9 PCs, p=0.2581, Mann-Whitney test) of SOM-mediated IPSCs in CA1 PCs from STD and EE mice. **c)** Representative traces of two consecutive SOM-mediated IPSCs at 20 Hz in CA1 PCs from STD (black) and EE (red) mice.

**Supplementary Figure 5:**
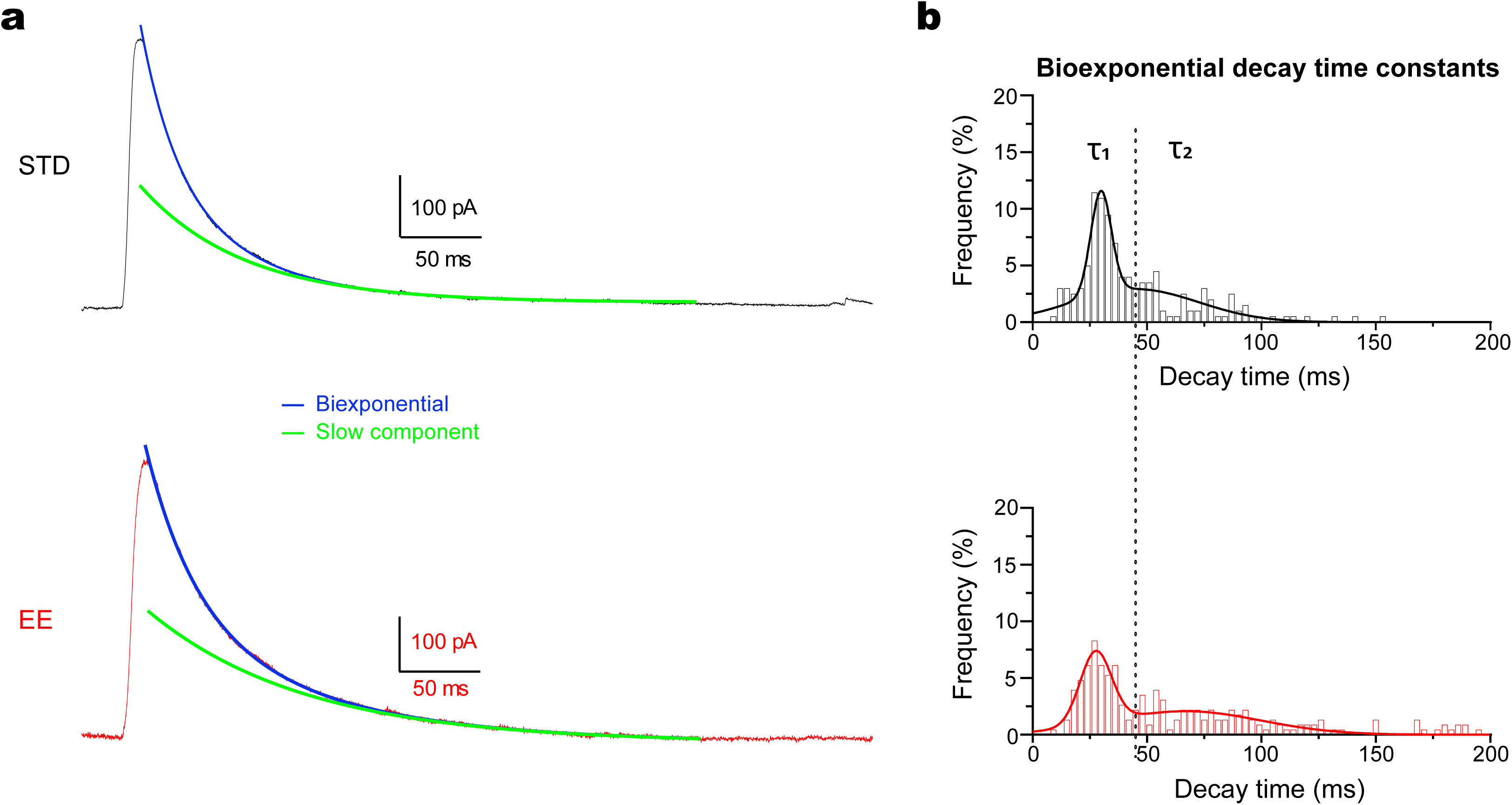
Biexponential decay time course of SOM-mediated IPSCs. **a)** Representative average traces of SOM-mediated IPSCs in CA1 PCs evoked by light stimulation of SOM interneurons for STD (black) and EE (red) mice. Blue lines indicate biexponential fits and green lines indicate the isolated slow decay component. The slow component in example traces was prolonged in EE (STD: τ_1_ = 28.8 ms and τ_2_ = 65.4 ms vs EE: τ_1_ = 31.4 ms and τ_2_ = 99.6 ms) **b)** Frequency distribution of the decay time constants after fitting a biexponential curve to the SOM-mediated IPSC for STD (black) and EE (red) mice, illustrating the separation of fast (τ1) and slow (τ2) decay components and the rightward shift of the slow component in EE.

**Supplementary Figure 6:**
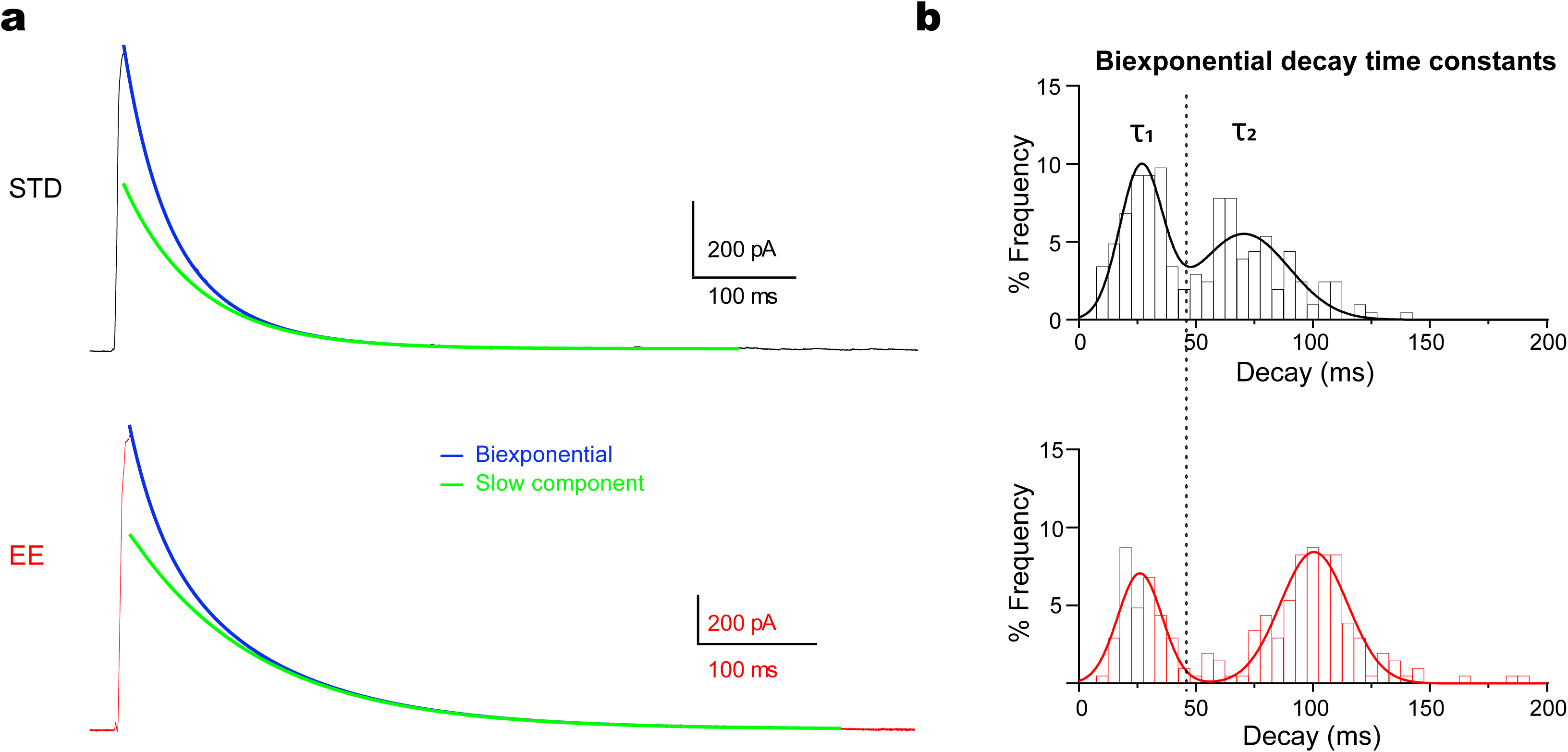
Biexponential decay time course of feedback IPSCs. **a)** Representative average traces of feedback IPSCs in CA1 PCs evoked by light stimulation of adjacent CA1 PCs for STD (black) and EE (red) mice. Blue lines indicate biexponential fits and green lines indicate the isolated slow decay component. The slow component in example traces was prolonged in EE (STD: τ_1_ = 33 ms and τ_2_ = 72.5 ms vs EE: τ_1_ = 31.4 ms and τ_2_ = 108.8 ms) **b)** Frequency distribution of the decay time constants after fitting a biexponential curve to the feedback IPSC for STD (black) and EE (red) mice, illustrating the separation of fast (τ1) and slow (τ2) decay components and the rightward shift of the slow component in EE.

**Supplementary Figure 7:**
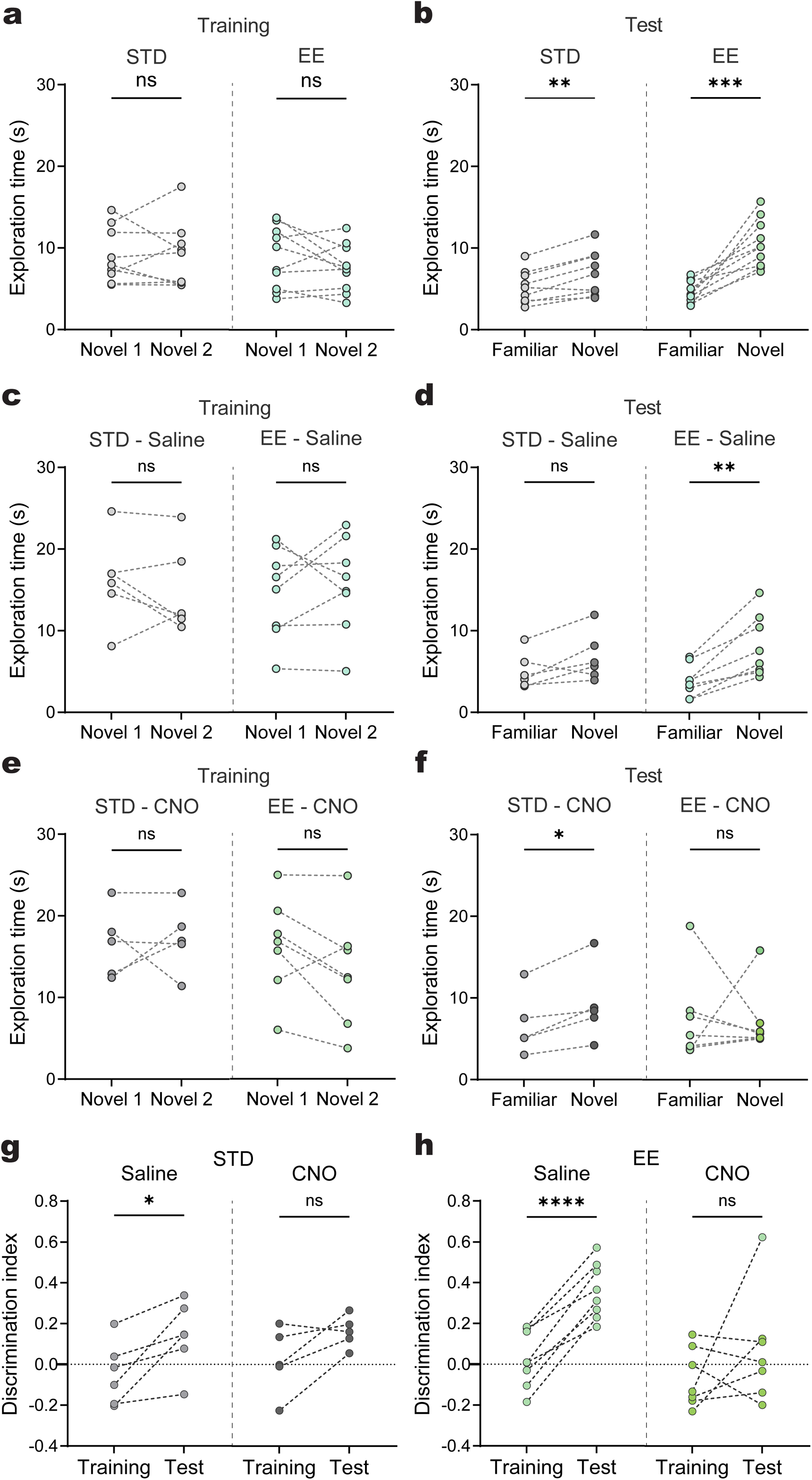
Object exploration during OLT training and test. **a)** Pairwise comparison of exploration time of the two novel objects during OLT (training) for STD (grey) and EE (green) mice (STD-Novel1 vs STD-Novel2: p=0.985, paired t-test, EE-Novel1 vs EE-Novel2: p=0.2153, paired t-test; STD, N=9 vs EE, N=10 mice). **b)** Pairwise comparison of exploration time of the two objects at familiar and novel locations during OLT (test) for STD (grey) and EE (green) mice (STD-Familiar vs STD-Novel: p=0.0017, paired t-test, EE-Familiar vs EE-Novel: p=0.0003, paired t-test; STD, N=9 vs EE, N=10 mice). **c)** Pairwise comparison of exploration time of the two novel objects during OLT (training) for STD (grey) and EE (green) mice injected with saline (STD-Novel1 vs STD-Novel2: p=0.3844, paired t-test, EE-Novel1 vs EE-Novel2: p=0.6030, paired t-test; STD-Sal, N=6 vs EE-Sal, N=8 mice). **d)** Pairwise comparison of exploration time of the two objects at familiar and novel locations during OLT (test) for STD (grey) and EE (green) mice injected with saline (STD-Familiar vs STD-Novel: p=0.0939, paired t-test, EE-Familiar vs EE-Novel: p=0.0014, paired t-test; STD-Sal, N=6 vs EE-Sal, N=8 mice). **e)** Pairwise comparison of exploration time of the two novel objects during OLT (training) for STD (grey) and EE (green) mice injected with CNO (STD-Novel1 vs STD-Novel2: p=0.7795, paired t-test, EE-Novel1 vs EE-Novel2: p=0.0981, paired t-test; STD-CNO, N=5 vs EE-CNO, N=7 mice). **f)** Pairwise comparison of exploration time of the two objects at familiar and novel locations during OLT (test) for STD (grey) and EE (green) mice injected with CNO (STD-Familiar vs STD-Novel: p=0.0167, paired t-test, EE-Familiar vs EE-Novel: p=0.8125, Wilcoxon matched-pairs signed-rank test; STD-CNO, N=5 vs EE-CNO, N=7 mice). **g)** Pairwise comparison of discrimination index during training versus test for STD mice injected with saline or CNO (STD-Training-Sal vs STD-Test-Sal: p=0.0235, paired t-test; STD-Training-CNO vs STD-Test-CNO: p=0.0806, paired t-test). **h)** Pairwise comparison of discrimination index during training versus test for STD mice injected with saline or CNO (EE-Training-Sal vs EE-Test-Sal: p<0.0001, paired t-test; EE-Training-CNO vs EE-Test-CNO: p=0.3008, paired t-test). Data are shown as mean ± s.e.m. * p<0.05, ** p<0.01, *** p<0.001 and **** p<0.0001.

